# Rapid prey capture learning slowly re-sets activity set points in rodent binocular visual cortex

**DOI:** 10.1101/2025.04.11.648036

**Authors:** Daniel P. Leman, Brian A. Cary, Diane Bissen, Wei Wen, Brian J. Lane, M. Regis Shanley, Nicole F. Wong, Krish B. Bhut, Gina G. Turrigiano

## Abstract

Neurons within primary visual cortex (V1) possess stable firing rate set points to which they faithfully return when perturbed by passive sensory manipulations. Chronic recordings in vivo suggest that these set points are present early in postnatal development and are stable through adulthood, raising the possibility that once they are established, they are largely immutable. Here we challenge this idea, using an ethological, vision-dependent prey capture learning paradigm in juvenile (critical period) rats. Juvenile rats rapidly (in one day) became proficient at catching crickets. This learning was impaired when V1 was inactivated, and was accompanied by an increase in the number of binocular V1 (V1b) neurons with firing reliably tied to specific behavioral epochs during hunts, indicating that hunting drives rapid functional plasticity within V1b. Remarkably, this rapid learning set in motion a slow, state-dependent increase in V1b firing, that began after learning was complete and persisted for days. This upward firing rate plasticity was gradual, occurred selectively during wake states, and in L2/3 was driven by a TNFα-dependent increase in excitatory synapses and excitatory synaptic charge onto pyramidal neurons – all features of homeostatic forms of plasticity within V1. Finally, TNFα inhibition administered after learning was complete diminished this increase in excitability and impaired the retention of hunting skills. Taken together, these data suggest that naturalistic learning in juvenile animals co-opts homeostatic forms of plasticity to reset firing rate setpoints within V1b, in a process that facilitates skill consolidation.

## INTRODUCTION

Experience-dependent plasticity in young mammals is essential for the proper wiring and functional maturation of neocortical circuits (Espinosa and Stryker, 2012). This process of circuit refinement requires cooperation between Hebbian forms of plasticity that use correlated or decorrelated activity to strengthen or weaken specific connections, and homeostatic forms that regulate excitability to stabilize firing and other network features around set point values (Abbott and Nelson, 2000; Turrigiano and Nelson, 2004; Wen and Turrigiano, 2024). In primary visual cortex (V1), this interplay has been extensively studied using sensory deprivation paradigms, where closing one eye induces a drop in network activity that is then slowly restored to baseline, through the sequential induction of Hebbian and homeostatic forms of plasticity (Hengen et al., 2016, 2013; Kaneko et al., 2008; Keck et al., 2013; Mrsic-Flogel et al., 2007). Recent experiments in V1 have followed firing rates over time during such sensory manipulations, and have demonstrated that homeostatic mechanisms slowly restore mean firing rates at both the individual neuron and network levels (Hengen et al., 2016; Torrado Pacheco et al., 2021), suggesting that neurons have individual firing rate set points (or firing rate “zones”) to which they faithfully return after perturbations. These set points are already established by the classic V1 critical period, and appear to be stable well into late adulthood (Hengen et al., 2016, 2013; Torrado Pacheco et al., 2021; Barnes et al., 2015; Wu et al., 2020; McGregor et al., 2024; Dhawale et al., 2017; Jensen et al., 2022). An outstanding question is whether these firing rate setpoints – once established – are largely invariant, or if they can be modulated or reset by salient experiences.

Prey capture learning in rodents is a vision-dependent learned behavior, making it an ideal paradigm for testing the idea that homeostatic setpoints in V1 might be malleable. Rodents are opportunistic omnivores (Landry, 1970; Clark, 1982) and hunt insects for food in the wild (Polsky, 1978; Dean and Redgrave, 1984; Langley, 1989) and in the lab (Hoy et al., 2016, 2019; Zhao et al., 2019; Michaiel et al., 2020; Groves Kuhnle et al., 2022). While the drive to hunt is instinctive, rodents must practice to become proficient; in both adult and juvenile mice, the time to capture prey decreases by about an order of magnitude across repeated hunting sessions (Hoy et al., 2016; Groves Kuhnle et al., 2022; Galvin et al., 2021; Bissen et al., 2025), presumably due to sensorimotor plasticity that enhances the ability to detect, track, intersect, and grasp prey. Binocular vision is the primary sense that guides the detection and pursuit of prey in rodents (Hoy et al., 2016, 2019; Michaiel et al., 2020; Galvin et al., 2021; Johnson et al., 2021; Holmgren et al., 2021); and prey capture learning enhances spine density and dynamics within L5 of mouse binocular V1 (V1b) during the critical period (Bissen et al., 2025), but whether this learning is able to perturb firing rate setpoints within V1b is unknown.

To address this, we obtained chronic extracellular recordings from upper layers of V1b of juvenile (critical period) Long Evans rats before, during, and after they learned to hunt live crickets. Learning was rapid, was prevented by chemogenetic inhibition of V1, and was accompanied by a dramatic increase in the proportion of V1b neurons with firing strongly tied to pursuit behavior. Mean firing rates in V1b were stable prior to and immediately after learning but began to slowly increase a few hours later and roughly doubled by ∼60 hours post-hunting. This slow increase in network excitability was accompanied by a TNFα-dependent increase in excitatory synapses and excitatory charge onto L2/3 pyramidal neurons, and occurred primarily during active wake states, all signatures of upward excitatory homeostatic synaptic plasticity (Barnes et al., 2022, 2017; Hengen et al., 2016; Kaneko et al., 2008; Steinmetz and Turrigiano, 2010; Stellwagen and Malenka, 2006; Turrigiano, 2012). Finally, disrupting TNFα signaling post-learning reduced the excitability change in L2/3 pyramidal neurons, and impaired the retention of hunting skills. These findings suggest that ethologically relevant learning initiates a slow transition to a new V1 network setpoint, that is driven by homeostatic forms of plasticity and enhances skill retention. More broadly, our findings demonstrate that homeostatic activity setpoints are themselves substrates for experience-dependent plasticity.

## RESULTS

### Prey capture learning in juvenile rats is rapid and requires visual cortex

Prey capture learning is a complex behavior that involves the cooperation of many brain regions to enable the detection, tracking, and capture of moving prey, including visual circuits in the superior colliculus (Hoy et al., 2019; Shang et al., 2019). Whether V1 is also required for prey capture learning is less clear (Bissen et al., 2025; Zhao et al., 2019). To assess whether V1 is necessary, and whether vision-dependent learning can induce long-lasting changes in V1 activity, we designed a behavioral chamber that can be converted between “hunting” and “home cage” configurations to enable behavioral testing while allowing continuous, multiday video monitoring and electrophysiological recordings before, during, and after learning in critical period Long-Evans rats. The “Hunting” configuration (Figure 1A) features 4 Arduino-controlled dispensers that dispense individual live crickets in randomized order (STAR Methods) (Groves Kuhnle et al., 2022; Bissen et al., 2025). “Home cage” configuration (Figure 1B) includes bedding, nesting materials, food and water *ad libitum*, and an adjacently housed littermate for social interaction. We then developed a single-day learning paradigm consisting of 3 hunting sessions, comprised of 10 hunts each, with 1.5-hour rest periods between sessions (Figure 1C). Prior to hunting, rats underwent two days of habituation and were fed immobilized crickets to overcome neophobia (Groves Kuhnle et al., 2022). When first presented with live crickets, naïve rats were highly motivated to pursue but had difficulty successfully intercepting and capturing crickets (Figure 1D, top, Movie S1); however, they rapidly improved, and by the third session were much faster (Figure 1D, bottom, Movie S2). Quantification of capture times revealed a significant improvement across sessions (Figures 1D-E). There was no significant difference between males and females (Figure S1A), so data were combined, and both sexes were used for all subsequent experiments. To determine whether improved performance was due to faster initiation or shorter pursuits, we measured latency to attack (from release to pursuit initiation) and pursuit duration (from initiation to capture) for each hunt and found that both measures decreased significantly as learning progressed (Figures 1F-G).

**Figure 1.**
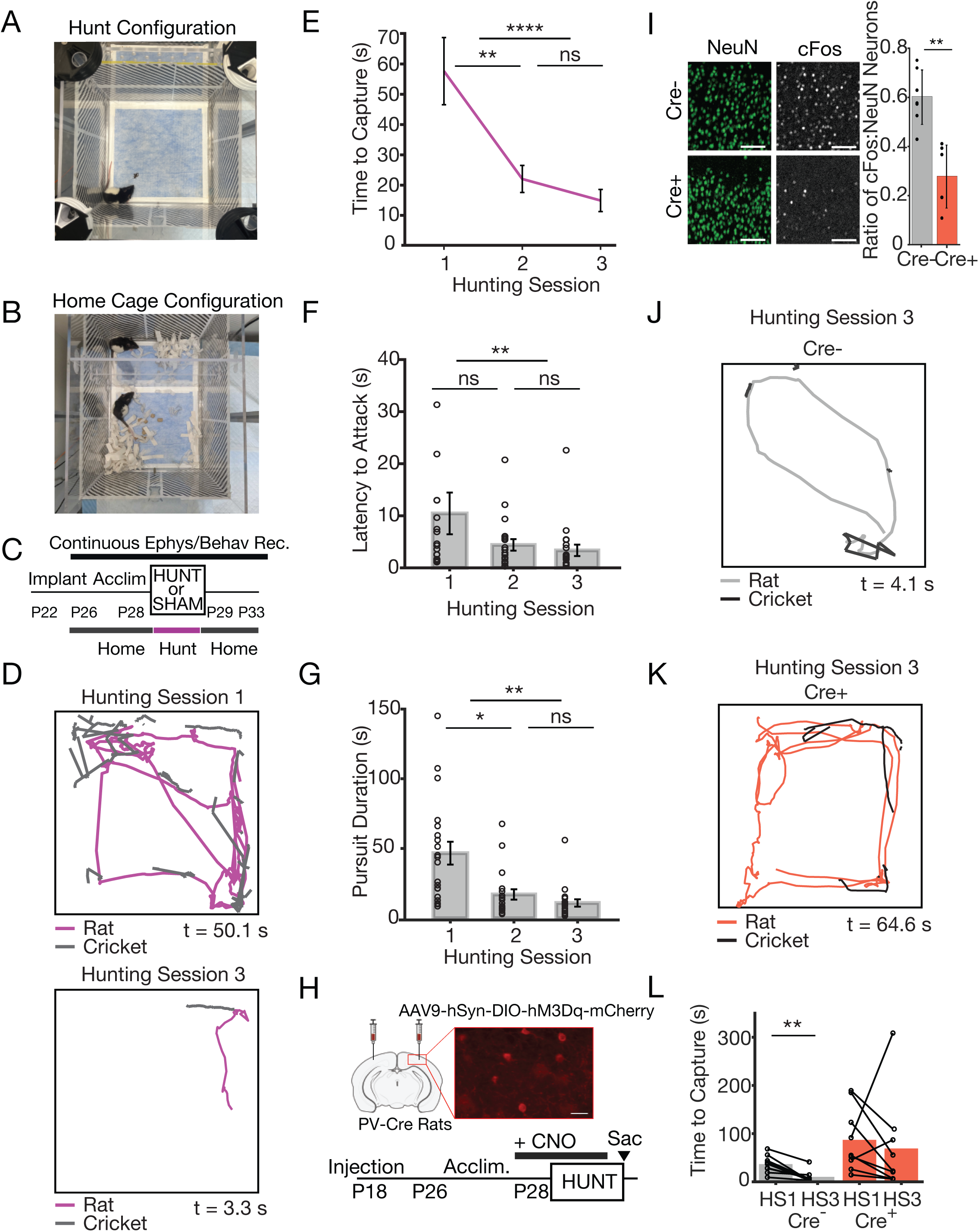
Rapid, V1-dependent prey capture learning in juvenile rats. **A.** Prey capture learning arena in “hunting” configuration with rat and cricket. **B.** Prey capture learning arena in “home cage” configuration with rat, bedding, food, water, and littermate in adjacent chamber for social interaction. **C.** Experimental timeline for chronic behavior and electrophysiology recordings for prey capture learning. **D.** Example trajectories of rat and cricket during one hunt by naïve (hunting session 1) or experienced (hunting session 3) rat. Magenta: rat position, black: cricket position; discontinuities due to cricket jumps. Time to capture is signified by t (seconds). **E-G.** (**E**) Average time to capture, (**F**) latency to attack, and (**G**) pursuit durations for electrode-implanted rats for hunting sessions 1-3. Circles represent individual animal averages per session (n=20 rats, 10 hunts/session) error bars represent the mean +/- SEM. Friedmann Test with Dunn Correction, Time to capture: HS1-HS2: p=0.003, HS1-HS3: p<0.0001, HS2-HS3: p=0.5; Latency to Attack: HS1-HS2: p=0.05, HS1-HS3: p=0.003, HS2-HS3: p>0.9; Pursuit Duration: HS1-HS2: p=0.006, HS1-HS3: p<0.0001, HS2-HS3: p=0.4. **H.** Experimental timeline for DREADDs-mediated chemogenetic inhibition of V1 during prey capture learning. Inset: PV interneurons expressing excitatory DREADDs (labeled with mCherry). Scale bar=50μm. **I.** Fraction of NeuN+ pyramidal neurons that are also cFos+, in littermates with (Cre+) or without (Cre-) excitatory DREADDs expressed in PV neurons. Wilcoxon Rank Sum test, p=0.001. PV-Cre-: n=7 rats, cFos: n=1955 neurons, NeuN: n=3332 neurons; PV-Cre+, n=6 rats, cFos: n=855 neurons, NeuN: n=2996 neurons. **J, K.** Example trajectories of rat and cricket from hunting session 3 for (**J**) Cre- and (**K**) Cre+ animal in presence of CNO. **L.** Average time to capture for hunting session 1 vs 3 on D1 per animal for PV-Cre- vs PV-Cre+ rats (Cre-: n=9; Cre+: n=9), Wilcoxon Sign Rank test, p=0.004. Here and onward, all error bars represent the mean +/- SEM unless otherwise indicated.

Vision is the dominant sensory modality used during prey capture in rodents (Hoy et al., 2016, 2019; Zhao et al., 2019; Michaiel et al., 2020; Galvin et al., 2021; Bissen et al., 2025; Johnson et al., 2021; Holmgren et al., 2021; Procacci et al., 2020), but the role of V1 is less clear. To determine whether V1 is necessary for learning, we inactivated V1 using viral expression of a Cre-recombinase (Cre)-dependent excitatory DREADDs construct (AAV9-hSyn-DIO-hM3Dq-mCherry) in V1, targeted to Parvalbumin-positive (PV) interneurons using transgenic PV-Cre rats (Yu et al., 2018) (Figure 1H). Clozapine-N-Oxide (CNO, 0.05mg/ml) was administered to rats via drinking water (Wen and Turrigiano, 2021; Bottorff et al., 2024) for approximately 16 hours prior to hunting session 1 and between subsequent sessions; control animals were Cre^-^ littermates that received identical virus injection and CNO administration. To confirm this paradigm reduced V1 activity, we sacrificed rats following hunting session 3 and immunolabeled V1 slices for NeuN (to label pyramidal neurons) (Bottorff et al., 2024; Chattopadhyaya et al., 2004; Nahmani and Turrigiano, 2014) and the immediate early gene cFos. We observed a significant decrease in the fraction of NeuN^+^ neurons that were also cFos^+^ in Cre^+^ compared to Cre^-^ littermates (Figure 1I), indicating a robust suppression of V1 activity. Next, we compared hunting performance; Cre^-^ animals decreased time to capture across sessions as expected, and by session 3 rapidly caught prey (Figures 1J, 1L, S1B, and Movie S3). In contrast, while Cre^+^ rats readily chased crickets (Figure 1K, Movie S4), they took much longer to catch prey and failed to improve across sessions (Figures 1K,L, and S1B). Thus, V1 plays a critical role in mediating this visually-guided behavior and is required for rapid prey capture learning.

### The activity of V1b neurons becomes tuned to behavior during prey capture learning

To determine whether vision-dependent naturalistic learning drives changes in the activity of V1 neurons, we recorded single-unit activity in layers 2/3 and 4 (L2/3, L4) of V1b continuously for 7 days in freely behaving juvenile rats, as described previously (Hengen et al., 2016; Torrado Pacheco et al., 2021; Bottorff et al., 2024). We targeted V1b because rodents use binocular vision to guide predation (Hoy et al., 2016, 2019; Michaiel et al., 2020; Galvin et al., 2021; Johnson et al., 2021; Holmgren et al., 2021), and upper layers because of extensive information on homeostatic mechanisms and firing rate homeostasis in L2/3 and 4 (Hengen et al., 2016, 2013; Keck et al., 2013; Torrado Pacheco et al., 2021; Wen and Turrigiano, 2021; Bottorff et al., 2024; Lambo and Turrigiano, 2013; Maffei and Turrigiano, 2008; Keck et al., 2011; Gainey and Feldman, 2017). Recordings were initiated during habituation (P26 or 27), maintained during learning (P28 or 29), and continued for 66 hours following the last hunting session (Figure 1C). We isolated regular spiking (RSU, predominantly excitatory neurons) and fast-spiking (FSU, putative inhibitory interneurons) single units as previously described (Hengen et al., 2016, 2013; Torrado Pacheco et al., 2021; Bottorff et al., 2024; Niell and Stryker, 2008; Cardin et al., 2007) (STAR Methods). Control animals underwent the same paradigm as hunting counterparts (including receiving immobilized crickets during habituation), except in “sham” hunting sessions the cricket dispensers were empty, and rats were given a food pellet at the end of each session.

To assess if V1b activity becomes tied to specific behavioral epochs during learning, we isolated individual RSUs that we could follow across all three prey capture learning sessions. We segmented each hunt into 4 epochs: inter-cricket interval (“ICI”, from consumption end to next cricket release), latency to attack (time from cricket release to pursuit), pursuit duration (start of pursuit to capture), and consumption (from capture to the end of eating) (Figures 2A and 2B). Initially, most RSU firing rates were not significantly modulated by hunting (Figure 2C, top), but as learning progressed, many RSUs developed robust responses to the pursuit epoch (Figure 2C, bottom, red arrows). To quantify this, we first calculated the percent change in firing for each neuron relative to the preceding ICI, and took the population average (117 neurons, 6 animals) during pursuit and consumption epochs across sessions. RSU firing increased slightly during pursuits in hunting session 1 (Figure 2D, left), and the magnitude of this modulation increased to ∼50% during sessions 2 and 3 (Figure 2D, left). In contrast, RSU firing across the population was not significantly elevated during consumption for any session (Figure 2D, right).

**Figure 2.**
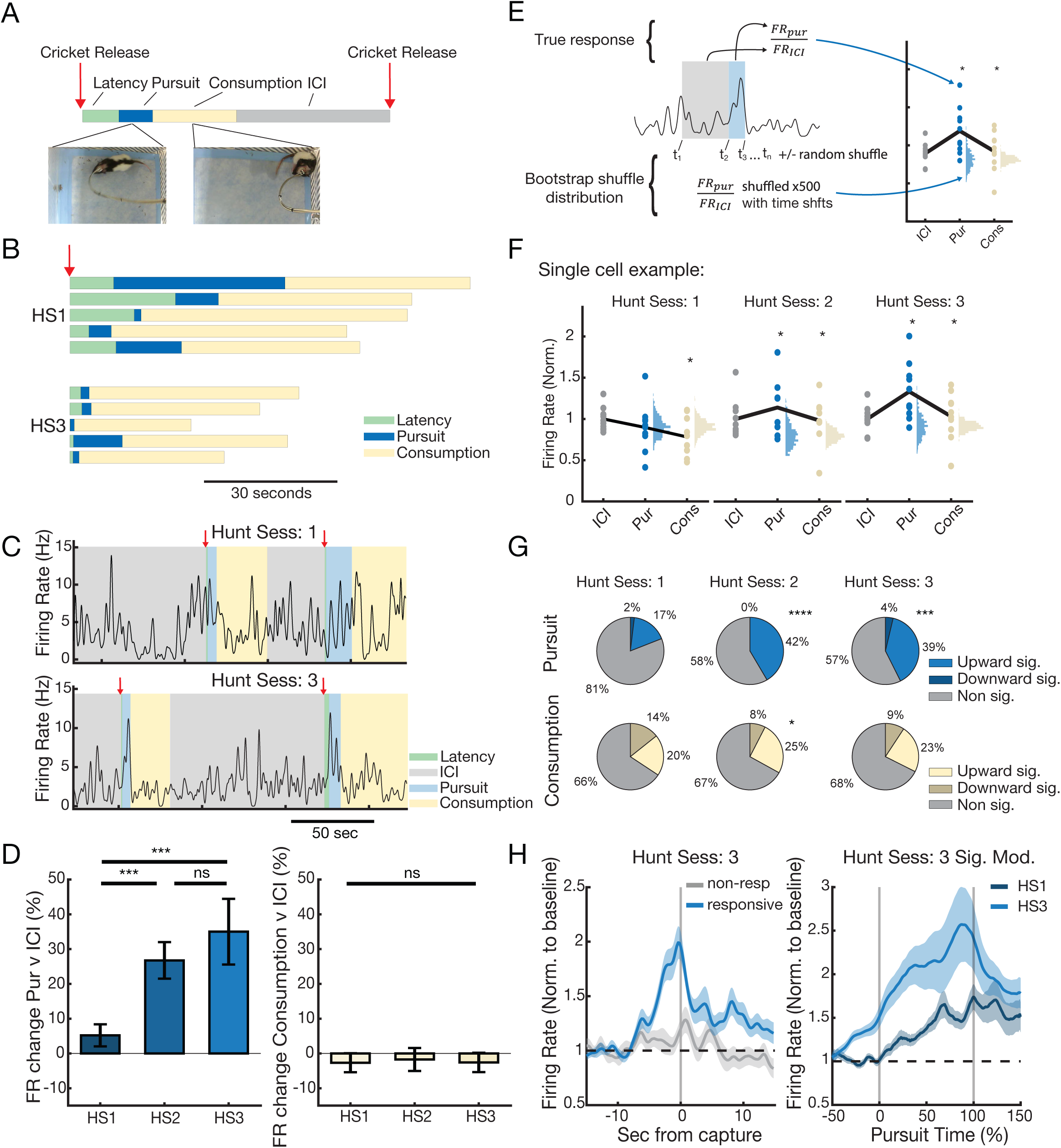
Firing of regular spiking V1b neurons becomes tuned to behavior during prey capture learning. **A.** Example ethogram showing behavioral sequence during a hunt. ICI = inter-cricket interval. **B.** 5 example ethograms from the first (HS1) and third (HS3) hunting session, illustrating variability in timing. **C.** Two example firing rate (FR) traces from the same cell on HS1 and HS3, respectively. Behavioral sequence labels indicated with colored boxes. **D.** Percentage change in FR from ICI during pursuit (left) and consumption (right) across hunting sessions (n=111 cells from 6 animals). Repeated measures ANOVA (rmANOVA) with Tukey-Kramer post hoc: Pursuit: HS1-HS2: p=0.0004, HS1-HS3: p=0.006, HS2-HS3: p=0.5. Consumption: HS1-HS2: p>0.9, HS1-HS3: p>0.9, HS2-HS3: p>0.9. **E.** Schematic showing bootstrap approach to determining significant FR modulation during hunting behaviors. **F.** Illustration of bootstrap results showing a single cell’s responses across the 3 hunting sessions. Circles indicate actual normalized FRs during given behavior. Sideways histograms represent calculated bootstrap shuffle distributions. Asterisks indicate bootstrap significance (<5^th^ or >95^th^ percentiles). **G.** Percentage breakdown of up/down/non-significant modulation of the RSUs across prey capture learning (n=103 cells from 6 animals). Friedman Test with Tukey-Kramer post hoc; Pursuit (top row): HS1-HS2: p<0.0001, HS1-HS3: p=0.0007. Consumption (bottom row): HS1-HS2: p=0.04, HS1-HS3: p=0.2. **H.** Left, normalized FR aligned to capture for responsive and non-responsive RSUs on hunting session 3. Right, normalized FR against normalized pursuit time for cells that are pursuit responsive on HS3, comparing HS1 and HS3 responses. FRs are normalized to an 8 second window preceding capture events (left) or pursuit initiation (right).

To determine what fraction of neurons became responsive during pursuit epochs, we compared firing rates during pursuit (normalized to ICI as above) to a bootstrap shuffled distribution for each neuron for each hunting session (Figure 2E, STAR Methods). Many neurons that were initially unresponsive became significantly modulated by pursuit as learning progressed; a single neuron example is shown in Figure 2F. Across the population, there was a progressive increase in the number of significantly pursuit-modulated RSUs across hunting sessions, from 17% in session 1 to 39% by session 3 for positively modulated RSUs, while the small number of negatively modulated RSUs was unchanged (Figure 2G, top). In contrast, there was only a modest increase in RSUs modulated by consumption (Figure 2G, bottom). To examine the time course of pursuit-modulated firing, we aligned firing to capture during HS3, revealing higher activity in the responsive compared to the nonresponsive population for several seconds before and after capture (Figure 2H, left). We next compared the firing of neurons that were pursuit active in HS3 to their activity in HS1; because pursuit time shortens across sessions, we scaled each response on the time axis to align firing to start (0%) and end (100%) of pursuit. This analysis revealed larger and more sustained increases in firing in HS3, that peaked around time of capture (Figure 2H, right).

In V1 of head-fixed mice firing rates are strongly modulated by movement speed (Niell and Stryker, 2010; Fu et al., 2014; Vinck et al., 2015; Mineault et al., 2016; Dadarlat and Stryker, 2017; Parker et al., 2020); to our knowledge only one study has examined speed modulation in freely moving rats where it is more modest (Guitchounts et al., 2020, but see Haggerty and Ji, 2015 for movement modulation). We thus wondered whether faster movement as animals learn to hunt could explain enhanced firing during pursuit epochs. Mean pursuit speed increased across hunting sessions from about 2 to about 3 cm/s (Figure S2A) and distance traveled decreased (Figures S2C), but maximum speed was similar across sessions (7-8 cm/s, Figure S2B). Animal speeds aligned around capture showed similar dynamics, but with a mild increase right before capture in session 3 (Figure S2D). To determine if the increase in mean pursuit speed between HS1 and HS3 could contribute to enhanced firing, we quantified firing rate as a function of speed on the hunting day (Figure S2E); while speed did modulate firing, this effect was relatively modest and saturated at speeds <1 cm/s, well below the mean pursuit speed (Figure S2E, range denoted by grey shaded region). Thus, differences in speed alone cannot account for enhanced V1b responses during pursuit. Taken together, these data demonstrate that prey capture learning amplifies V1b responses during specific behavioral epochs.

### Prey capture learning induces a slowly developing and persistent increase in V1b firing rates

We next investigated whether rapid vision-dependent learning has long-lasting effects on activity setpoints in V1b. We first compared RSU firing rate distributions at baseline and several days after learning (60 hours post-hunting). As expected, there was no significant difference between baseline distributions from hunt and sham animals (Figure 3A). Surprisingly, at 60 hours post-hunting, the hunt distribution shifted significantly toward higher firing rates relative to the sham condition (Figure 3B). This shift toward higher firing rates was also apparent when comparing baseline to 60 hours post-hunt for hunt animals and was most prominent at the low end of the distribution (Figure 3C).

**Figure 3.**
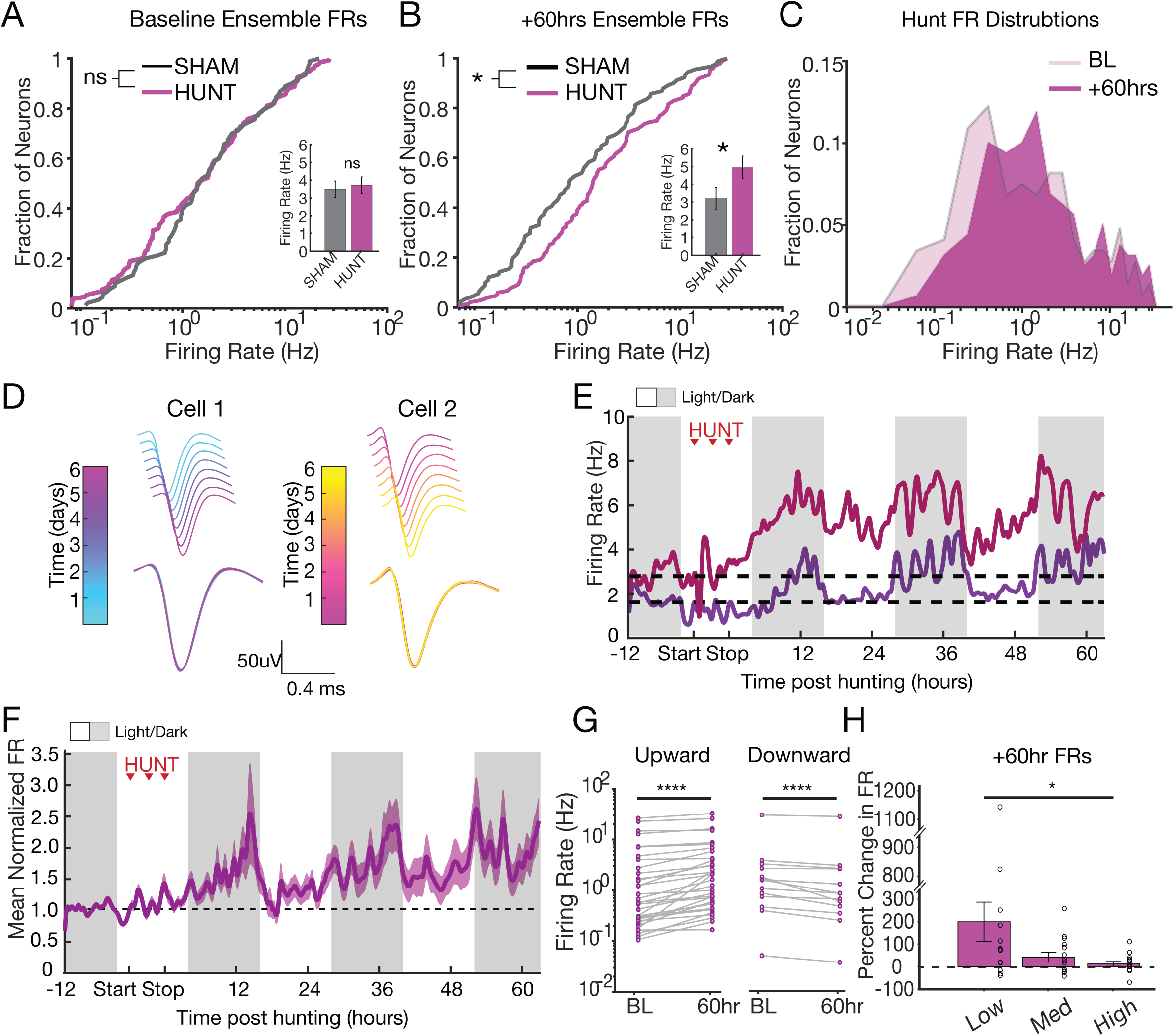
Prey capture learning induces a slowly developing and persistent increase in V1b firing rates. **A.** Distributions of ensemble FRs for Sham (grey) and Hunt (magenta) conditions at baseline. SHAM: n=109 neurons, Hunt: n=140 neurons. Anderson Darling test, p=0.2. Inset: Average ensemble firing rates. Wilcoxon Rank Sum, p=0.09. **B.** Distributions of ensemble FRs for Sham (grey) and Hunt (magenta) conditions following prey capture learning or sham. SHAM: n=91 neurons, Hunt: n=152 neurons, Anderson Darling test, p=0.01, Inset: Average ensemble firing rates. Wilcoxon Rank Sum, p=0.01. **C.** Distribution of firing rates from the hunt condition comparing baseline (BL) and 60 hours post-learning (+60hrs). **D.** Average waveform traces for two example RSUs recorded from the same electrode. Each spike waveform was color-coded by day to illustrate stability; the overlay across days is shown below. **E.** FRs of two individual RSUs (red and purple) measured continuously for 12 hours prior to and 62 hours after prey capture learning. Red arrows here and onward indicate individual hunting sessions; white/grey bars indicate light/dark periods. **F.** Mean, normalized FRs of all continuously recorded RSUs (n=45 from 6 animals). Here and onward, shaded area around mean firing rate indicates SEM.**G.** Average FRs for continuously recorded RSUs in Figure 3F, with increased (left, n=29) and decreased (right, n=16) firing rates from baseline, measured at 60 hours post-prey capture learning. Wilcoxon Sign Rank test, p<0.001 and p<0.001. Purple line indicates mean firing rates (baseline: 2.9Hz, 60hrs: 3.6Hz). **H.** Percent change in RSU firing rates 60 hours post-hunting, grouped into tertiles (low, medium, high) based on baseline firing rates. One-way ANOVA, p=0.04.

To determine the time course of this shift, we followed individual RSUs as described previously (Hengen et al., 2016; Torrado Pacheco et al., 2021; Bottorff et al., 2024) by spike-sorting on our entire continuously-recorded dataset and identifying neurons that we could reliably track throughout the experiment (Figures 3D-E). Two examples of continuously recorded neurons are shown in Figure 3E; both underwent a slow increase in firing that commenced several hours after hunting session 3 and remained elevated well-above baseline by 60 hours post-hunting. The same pattern was seen in the normalized RSU population average for continuously recorded neurons, where firing slowly increased, with a pronounced light/dark oscillation, until roughly doubling by 50-60 hours post-hunting (Figures 3F-G; 45 neurons from 6 animals). Across the population there was a significant increase in firing (Figure S3A, B), with about 2/3 of neurons increasing (Figure 3G, Upward) and 1/3 decreasing (Figure 3G, Downward). To determine whether this behavior depends on initial firing rates, we divided RSUs into terciles by baseline firing rates and plotted percent change at 60 hours post-hunting. Low firing rate neurons underwent the largest change, while high firing rate neurons changed the least (Figure 3H), consistent with our findings that the firing rate distribution increased most at the low end (Figure 3C). In contrast to the dramatic, long-lasting changes in hunt firing rates, sham hunting produced only a transient change that returned to baseline by 60 hours (19 neurons from 6 animals, Figure S3C). Surprisingly, neurons recorded from monocular V1 (V1m) also showed no significant change in firing following hunting (12 neurons from 4 animals, Figure S3D), suggesting this effect is confined to V1b. This slow change in firing is not driven by changes in movement speed after learning: average speed increased transiently on the day of hunting, but returned to a stable baseline immediately after (Figure S3E), and the modulation of firing rate by speed did not increase from baseline to +52 post-hunting, when firing rates are elevated (Figure S3F). Taken together, these data demonstrate that prey capture learning initiates a gradually developing and long-lasting perturbation of V1b mean excitatory firing rates, that manifests both at the population level and in the activity of individual neurons.

### Fast spiking firing rates also increase after prey capture learning

Increased RSU firing rates post-hunting could arise through a reduction in firing of inhibitory neurons. We therefore identified putative inhibitory fast-spiking units (FSU) based on spike waveform parameters (Figure 4A) (Hengen et al., 2016, 2013; Niell and Stryker, 2008; Cardin et al., 2007); STAR Methods) and analyzed their activity. Like RSUs, FSU firing rates (41 neurons from 6 animals) increased significantly during pursuit epochs as animals learned to hunt (Figure 4B, left), with no change during consumption (Figure 4B, right). The number of pursuit-responsive FSUs also increased significantly between HS1 and HS2 (Figure 4C), and like RSUs, the firing of this FSU population was elevated around capture relative to non-significantly modulated FSUs (Figure 4D, left), and showed a larger and more sustained increase in firing at HS3 relative to HS1, that peaked around time of capture (Figure 4D, right). Therefore, prey capture learning drives similar rapid changes in RSUs and FSUs, suggesting a wide-spread reconfiguration of V1b response properties.

**Figure 4.**
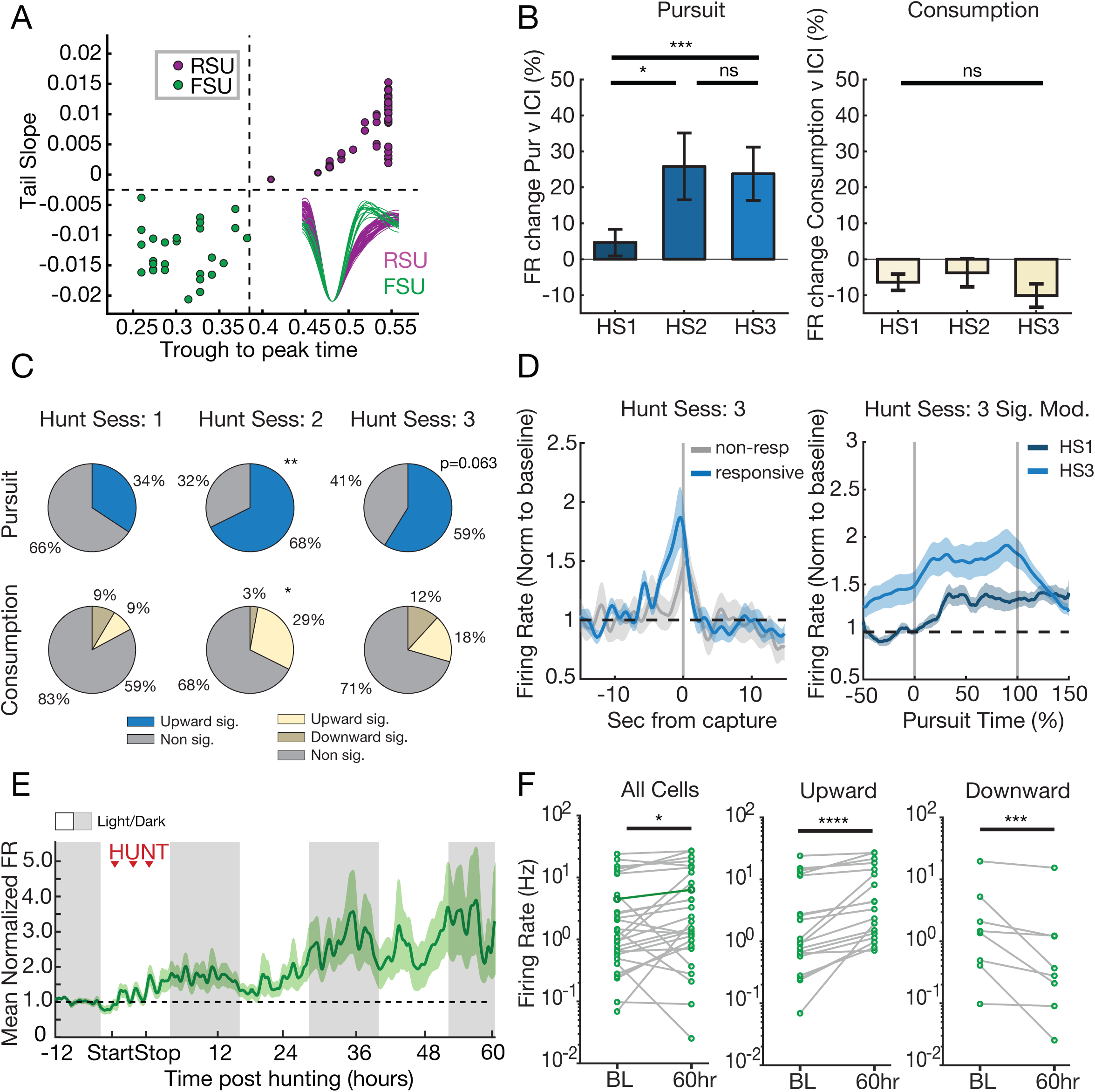
Fast-spiking firing rates are also persistently increased by prey capture learning. **A.** Plot of spike trough-to-peak vs. tail slope showing segregation into FSUs (green) and RSUs (magenta). **B.** Percentage change in FR during pursuit (left) and consumption (right) across hunting sessions (n=35 from 6 animals). Repeated measures ANOVA with Tukey-Kramer post hoc test, Pursuit: HS1-HS2: p=0.01, HS1-HS3: p=0.0007, HS2-HS3: p>0.9. Consumption rmANOVA: p=0.1. **C.** Pie charts indicating percentage breakdown of up/down/non-significant modulation of FSUs, determined by bootstrap analysis as in Figure 2G (n=34 from 6 animals). Friedman test with Tukey-Kramer post hoc correction, Pursuit (top row): HS1-HS2: p=0.006, HS1-HS3: p=0.06. Consumption (bottom row): HS1-HS2: p=0.03, HS1-HS3: p=0.8. **D.** Left, normalized FR aligned to capture for responsive and non-responsive FSUs on hunting session 3. Right, normalized FR against normalized pursuit time for cells that are pursuit responsive on HS3, comparing HS1 and HS3 responses. FRs are normalized as in Figure 2H. **E.** Mean, normalized FR of continuously recorded FSUs (n=28 from 6 animals). **F.** Left: mean FRs of individual FSUs measured at baseline (BL) and 60 hours post-prey capture learning (+60hr). Wilcoxon Sign Rank test, p=0.01. Middle: same, for FSUs with increased firing (n=20), Wilcoxon Sign Rank test, p<0.0001. Right: same, for FSUs with decreased firing (n=8), Wilcoxon Sign Rank test, p=0.008.

Next, we investigated whether prey capture learning triggers long-term changes in FSU firing rates, as it does for RSUs. For FSUs we could follow continuously (28 neurons from 6 animals), we observed a slowly developing increase in the mean normalized firing rate that persisted for at least 60 hours post hunting (Figure 4E). Like RSUs, comparing baseline to 60-hour post-hunt revealed a significant increase in FSU population firing rates (Figure 4F, left), with approximately 2/3 increasing (Figure 4F, middle) and 1/3 decreasing (Figure 4F, right). Thus, reduced FSU firing cannot account for increased RSU firing; instead, prey capture learning produces gradual, long-lasting changes in the activity of both of these excitatory and inhibitory circuit elements.

### Upward regulation of FR post-learning is gated by sleep/wake states

This gradual increase in network excitability (Figures 3F and 4E) is reminiscent of a slow homeostatic process, raising the interesting possibility that learning may rapidly “reset” activity setpoints within V1b, and subsequently set in motion a slow transition to these new setpoints. Further, sensory-driven firing rate homeostasis is gated by sleep/wake states, and the strong light/dark oscillation in post-hunting firing rate increases (Figures 3F and 4E) suggests this process could be similarly gated, since juvenile rats sleep more in the light and less in the dark (Cary and Turrigiano, 2021). To test this, we classified behavioral states of each animal into active wake (AW), quiet wake (QW), rapid eye movement (REM) sleep, and non-REM (NREM) sleep, using polysomnography as described (Hengen et al., 2016; Torrado Pacheco et al., 2021; Cary and Turrigiano, 2021) (STAR Methods). Because longer periods of sleep or wake drive stronger firing rate homeostasis (Hengen et al., 2016; Torrado Pacheco et al., 2021), we identified extended episodes of sleep and wake, and analyzed firing rate changes within those episodes (>30 mins in state; Miyawaki and Diba, 2016; Torrado Pacheco et al., 2021); REM episodes were excluded because they are generally short and there were too few extended episodes to reliably analyze.

In Figure 5A, the ensemble average firing rates from one post-hunt animal is shown color-coded by behavioral state; during wake, there was a clear increase in firing, while during sleep, there was a subtle decrease. To quantify this across animals, we calculated the z-scored change in firing across sleep/wake episodes for each state (AW, QW, NREM). In hunt animals, the largest increase was during AW, with a smaller increase in QW and a small decrease in NREM (Figure 5B, left). In contrast, there was no significant increase during wake states in sham animals, and a small decrease during NREM (Figure 5B, right), consistent with the subtler, transient change in network activity in this condition (Figure S3C). In addition, for hunt (Figure 5C), but not sham (Figure 5D) animals, the longer they spent in a state, the larger the change. There was no difference in the proportion of time spent in individual behavioral states between hunt and sham conditions, in either light (Figure S4A, top) or dark (Figure S4A, bottom) periods. Thus, the post-learning increase in firing rates is specific to wake states and largest in AW, and the magnitude of change correlates with time spent in that behavioral state.

**Figure 5.**
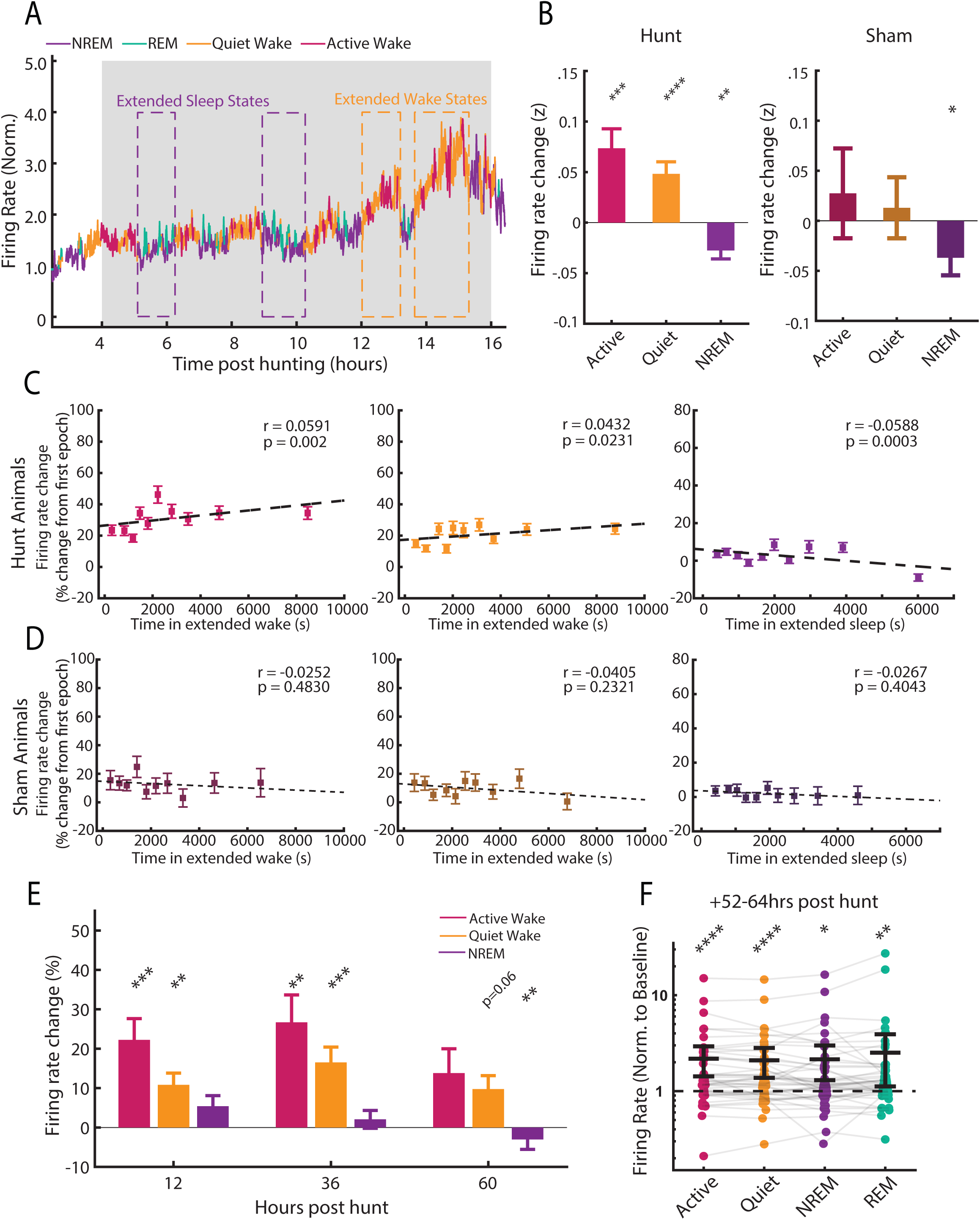
Learning-induced firing rate plasticity is gated by sleep/wake states. **A.** Ensemble average FR from one rat after hunting, color-coded by behavioral state. Two examples each of extended sleep and wake periods are highlighted. **B.** Mean change in z-scored FR across all extended quite wake, active wake, and NREM episodes after hunting (left) or sham (right). One-sided Wilcoxon Sign Rank test with Bonferroni Correction; p-values for Hunt condition (left): AW p=0.0006, QW p<0.0001, NREM p=0.0003. Sham (right): AW p>0.9, QW p>0.9, NREM p=0.045. Hunt: n=98 wake episodes; n=118 sleep episodes. Sham: n=60 wake episodes; n=52 sleep episodes. **C.** Correlation between percentage change in FR in active wake (left), quiet wake (middle), and NREM (right) and time from start of extended wake/sleep episodes, respectively. Change is calculated from FR in the first quiet wake epoch for wake episodes and from the first NREM epoch for sleep. Scatter data points are grouped into 10 equal sized groups with means +/- SEM for visualization. Pearson r and p values were computed on the ungrouped data: AW p=0.002, QW p=0.02, NREM p=0.0003. n=2761 QW data points, 2688 AW data points, 3861 NREM data points. **D.** Same as in (**C**) but for Sham animals. Pearson results: AW p=0.5, QW p=0.2, NREM p=0.4. n=871 QW data points, 772 AW data points, 975 NREM data points. **E.** FR change across extended wake/sleep episodes measured within Active/Quiet wake and NREM states in hunt animals. Sign rank test with Bonferroni corrected p-values: 12hr AW p=0.0001, QW p=0.003, NREM p>0.9; 36hr AW p=0.001, QW p=0.0003, NREM p=0.2; 60hr AW p=0.6, QW p=0.06, NREM p=0.003. **F.** Individual mean RSU firing rates post-hunt for each state, normalized to within-state firing rates in the baseline period. Sign rank test with Bonferroni corrected p-values: AW p=0.0004, QW p=0.001, NREM p=0.01, REM p=0.002. n=45 cells, 6 animals.

To determine whether the gradual stabilization of firing rates was reflected in the temporal profile of behavioral state-driven firing rate plasticity, we separately analyzed extended states at early (12hr), middle (36hr), and late (60hr) post-hunting timepoints. The largest percentage increase in firing during wake occurred in the middle timepoints, and by the late timepoint, wake no longer produced a significant increase and was balanced by a decrease during NREM (Figure 5E); results were similar when we analyzed z-scored firing rate (Figure S4B). Finally, to verify that wake-gated increases in firing were sustained regardless of subsequent behavioral state, we computed the change in firing rates at the end of the experiment (52-64 hours post-hunt) separately for each state and found similar increases across behavioral states (Figure 5F). In sum, after learning, RSU firing rates increase incrementally across many periods of active wake. Because upward firing rate homeostasis is also gated by sleep/wake states and is strongest during active wake (Hengen et al., 2016; Bottorff et al., 2024) these data support the idea that this slow increase in network excitability is driven by homeostatic forms of plasticity.

### Prey capture learning drives a TNFα-dependent increase in synapse number onto L2/3 pyramidal neurons

Homeostatic forms of plasticity in neocortical neurons include changes in excitatory synaptic strength and/or number (Keck et al., 2013; Torrado Pacheco et al., 2021; Turrigiano, 2012; Wen and Turrigiano, 2021; Lambo and Turrigiano, 2013; Turrigiano et al., 1998; Wierenga et al., 2006; Guerrero and Turrigiano, 2025), and are dependent on TNFα signaling (Barnes et al., 2017; Kaneko et al., 2008; Stellwagen and Malenka, 2006; Steinmetz and Turrigiano, 2010; Smilovic et al., 2022; Wen et al., 2025). Prey capture learning enhances spine density in the output layer (L5) of V1b (Bissen et al., 2025), but whether similar changes occur in the upper layers where they could contribute to the enhanced network activity we measure here is unknown. To quantify excitatory synapse number onto L2/3 pyramidal neurons, we quantified dendritic spine density as a proxy for excitatory synapses. We utilized a Thy1-YFP mouse line with sparse labeling in V1b L2/3 pyramidal neurons, which allowed for high-resolution reconstruction of entire dendritic arbors and accompanying spines (Figure 6A-C). Juvenile mice learn to hunt more slowly than rats (Groves Kuhnle et al., 2022; Bissen et al., 2025), so we used a modified paradigm consisting of 1 sham or hunt session/day for 3 days, followed by an additional “probe” session on day 6 (Figure 6D top, see Bissen et al., 2025), then quantified spine density on days 1 and 6 on all apical and basal dendritic segments from 6 fully reconstructed neurons/condition. The cumulative distribution of spine density across all dendritic segments in hunt animals was not significantly different on day 1 (Figure 6E) but by day 6 there was a significant shift toward higher densities (Figure 6F), indicating that learning induced a slow, robust, and widespread increase in spine density. The mean neuronal spine density was also significantly increased by ∼25% (Figure 6F, inset). Interestingly, there was a small but significant decrease in the cumulative distribution of spine head diameters after hunting (Figure S5A), which was not significant when calculated by cell (Figure S5A, inset); this was accompanied by a small decrease in mEPSC amplitude onto L2/3 pyramidal neurons (Figures S6B-C). These data suggest that post-learning spine formation generates a new population of slightly smaller excitatory synapses onto L2/3 pyramidal neurons.

**Figure 6.**
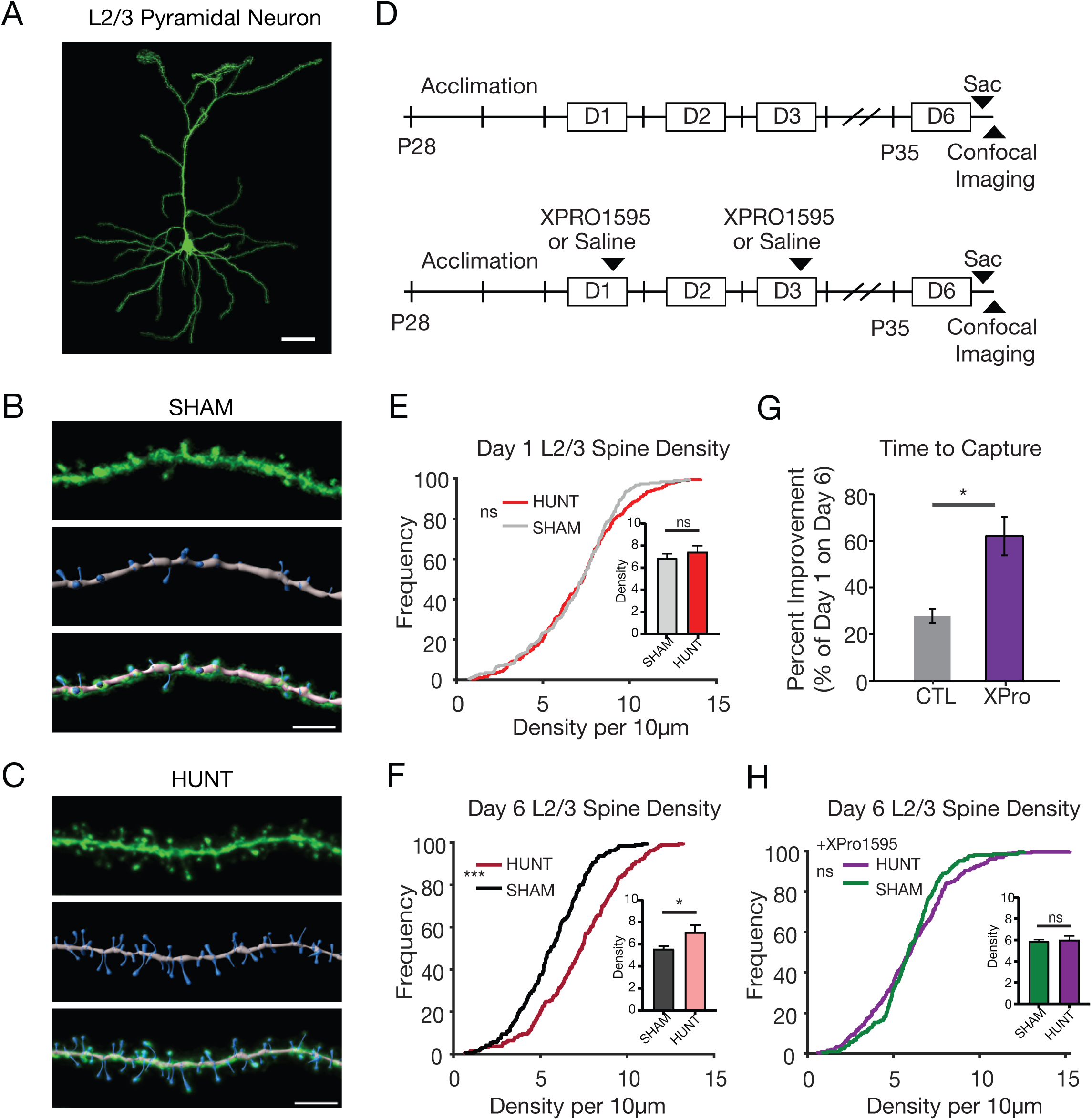
Learning drives a TNFα-dependent increase in L2/3 pyramidal neuron spine density. **A.** Example YFP-labeled mouse layer 2/3 V1b pyramidal neuron; scale bar = 50μm. **B.** Top: YFP-labeled dendritic segment from Sham condition; Middle: reconstruction of dendrites (grey) and spines (blue); Bottom: Overlay of top and middle images; scale bar = 5μm. **C.** Top: YFP-labeled dendritic segment from Hunt condition; Middle: reconstruction of dendrites (grey) and spines (blue); Bottom: Overlay of top and middle images. Scale bar = 5μm. **D. Top:** Experimental paradigm for mouse prey capture learning paradigm, consisting of 3 days of hunting (10 crickets per day, D1-D3), followed by a probe hunting session on D6. **Bottom:** Experimental paradigm used for XPRO1595 experiments; XPRO1595 was administered 1 hour following hunting sessions on Day 1 and again on Day 3. **E.** Cumulative distribution of L2/3 spine density, measured after Day 1 of prey capture learning. Sham: n= 197 spines, Hunt: n=235 spines from 5 sham and 6 hunting neurons, 3 animals per condition. Kolmogorov-Smirnov test, p=0.4. Inset: average spine density by neuron, Wilcoxon Sign Rank, p=0.4. **F.** Cumulative distribution of L2/3 spine density across all dendritic segments, measured 3 hours after learning was complete (Day 6). Sham: n=191 dendritic segments, Hunt: 189 dendritic segments, from 6 neurons and 3 animals per condition. Kolmogorov-Smirnov test, p<0.0001. Inset shows mean density by neuron; Unpaired t-test, p=0.046. **G.** Time to capture on Day 6, expressed as the percentage of Day 1 time to capture. CTL: n=8, XPro n=8, Unpaired t-test, p = 0.001. **H.** Cumulative distribution of L2/3 spine density in the XPro condition, measured as above. Sham: n=208 dendritic segments, Hunt: n=259 dendritic segments, from 6 neurons and 3 animals per condition. Kolmogorov-Smirnov test, p=0.1. Inset shows mean density by neuron; Unpaired t-test, p=0.8.

Homeostatic forms of plasticity, including sensory-deprivation induced spine plasticity in V1, are sensitive to TNFα signaling (Bissen et al., 2025; Barnes et al., 2022, 2017; Wen et al., 2025), as is spine addition in L5 pyramidal neurons following prey capture learning (Bissen et al., 2025), prompting us to ask if post-learning spine addition in L2/3 pyramidal neurons could be prevented by inhibiting this pathway. To answer this, we treated animals with XPRO1595 (XPro), a soluble TNFα scavenger that crosses the blood brain barrier and produces long-lasting inhibition of TNFR1 to block homeostatic plasticity (Bissen et al., 2025; Barnes et al., 2022, 2017; Wen et al., 2025). We injected XPRO1595 immediately post-hunt or sham on day 1, again 3 days later, and on day 6 analyzed dendritic spine density and diameter onto L2/3 pyramidal neurons from V1b (Figure S5B). Strikingly, treatment with XPro prevented the hunting-induced increase in spine density (Figures 6H); additionally, in the XPro condition, average spine density was similar to sham controls (inset, Figure H), suggesting XPro did not majorly impact baseline spine density. Finally, XPro treated mice had significantly less improvement in time to capture between day 1 and 6 than control mice (Figure 6G).

### Prey Capture Learning Shifts the E/I balance onto L2/3 Pyramidal Neurons to Favor Excitation

The slow increase in spine density post-learning suggests that increased RSU firing in L2/3 is driven at least in part by a net increase in excitatory drive. To determine if this is so, we obtained ex vivo whole-cell recordings from L2/3 pyramidal neurons from sham or hunt rats 3 days post-learning, and measured net excitatory and inhibitory synaptic charge in an “active slice” preparation (Figure 7A, Maffei et al., 2004; Trojanowski et al., 2021), by holding neurons successively at the experimentally-determined reversal potential for excitation and inhibition (Figure 7B, STAR Methods). Indeed, excitatory charge was significantly increased in the hunt condition (Figure 7C), and the amplitude and frequency of well-isolated excitatory currents were both significantly increased (Figures 7D-E). Further, these effects were prevented by treatment with XPro at the end of the last hunting session on day 1 (Figures 7C-E). In contrast, inhibition was unaffected: charge, amplitude, and frequency were not significantly different between conditions (Figures 7F-H). In a separate set of experiments we measured intrinsic excitability by generating firing rate vs. current (F-I) curves in the presence of synaptic blockers, which revealed overlapping F-I curves in the Sham and Hunt conditions (Figure S6D-E), and no significant difference in rheobase, input resistance, membrane potential, or area under the F-I curve (Figures S6F-I). Thus, changes in passive properties or intrinsic excitability do not contribute to the slow increase in firing of L2/3 neurons.

**Figure 7.**
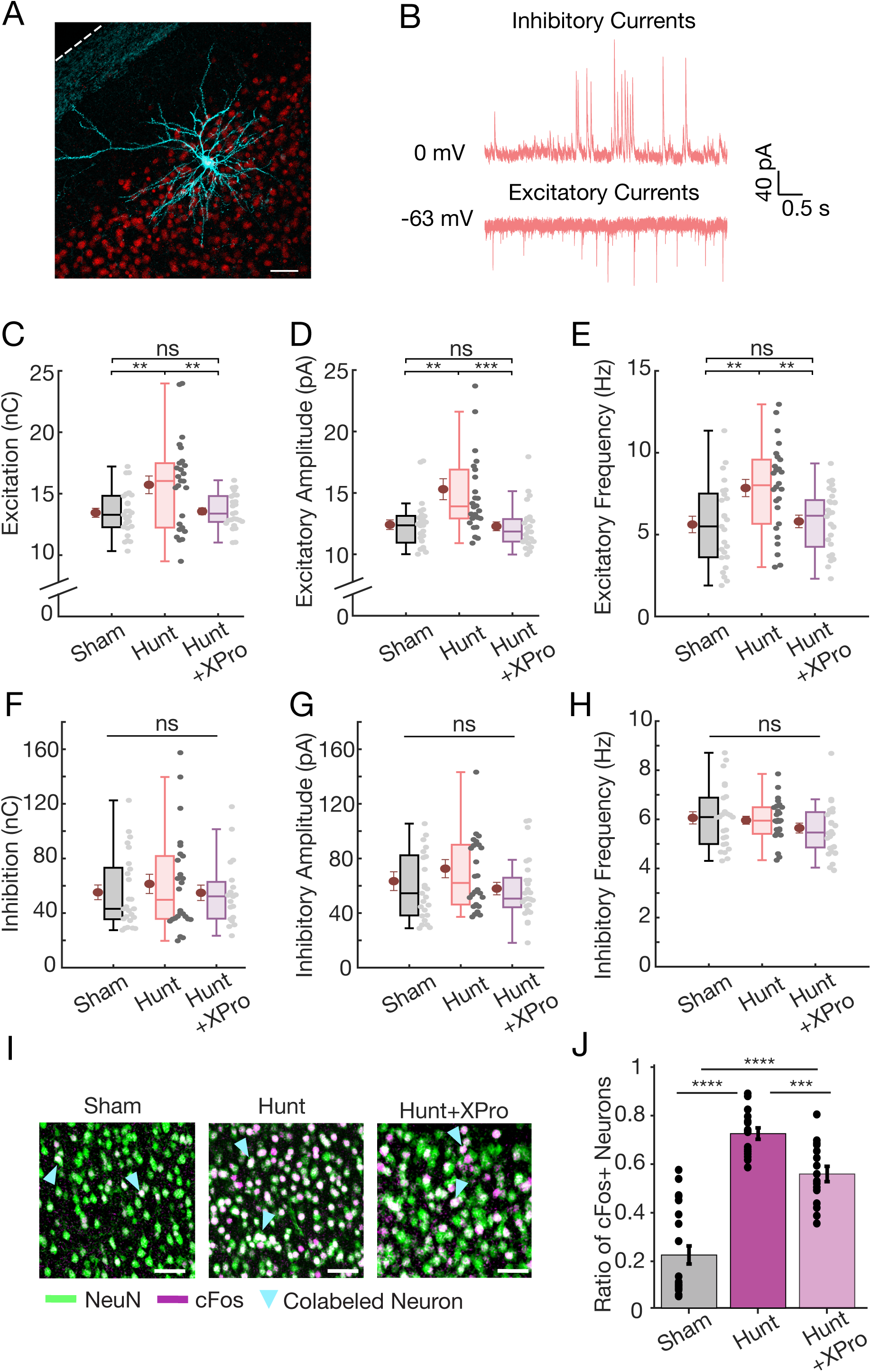
Prey capture learning increases excitatory but not inhibitory charge onto L2/3 pyramidal neurons. **A.** Representative image of a pyramidal neuron from upper L2/3 filled with biocytin during whole cell recording. Dashed line indicates pial surface; scale bar: 50 μm. **B.** Whole cell recording from V1b L2/3 pyramidal neuron measured in an “active” ex vivo slice preparation, 3 days post-learning. To isolate excitatory and inhibitory currents, neurons were held at the excitatory (upper example trace) or inhibitory (lower example trace) reversal potential in active ACSF. C. Comparison of the total excitatory synaptic charge for Sham, Hunt, or Hunt + XPro conditions. Here and throughout, each point is an individual neuron and red dots indicate mean ± S.E.M. One-way ANOVA with Tukey post-hoc correction: Sham vs. Hunt, p = 5.4E-3; Sham vs. XPro, p = 0.99; Hunt vs. XPro, p = 8.6E-3. **D.** Comparison of the mean amplitudes of well-isolated excitatory synaptic events. Kruskal-Wallis test with Tukey post-hoc correction: Sham vs. Hunt, p = 2.9E-3; Sham vs. XPro, p = 0.95; Hunt vs. XPro, p = 8.9E-4. **E.** Same as **C**, but for the mean frequencies. One-way ANOVA test with Tukey post-hoc correction: Sham vs. Hunt, p = 4.1E-3; Sham vs. Xpro, p = 0.96; Hunt vs. XPro, p = 9.3E-3. **F.** Comparison of the total inhibitory synaptic charge for the indicated conditions. Kruskal-Wallis test, p = 0.92. **G.** Comparison of the mean amplitudes of well-isolated inhibitory synaptic events. Kruskal-Wallis test, p = 0.29. **H.** Same as **F**, but for the mean frequencies. One-way ANOVA test, p = 0.34. **I.** Representative images of immunolabeled L2/3 V1 neurons for Sham, Hunt, and XPro conditions. Scale bar = 50μm. **J.** Fraction of NeuN+ V1 pyramidal neurons that are also cFos+ across each condition. One-way ANOVA with Bonferroni Correction: Sham vs. Hunt, p<0.0001, Sham vs. XPro, p>0.0001, Hunt vs. XPro, p=0.007; n=4 rats per condition.

Finally, we wanted to know if preventing increased excitatory drive onto L2/3 pyramidal neurons as above would reduce L2/3 pyramidal neuron activity in vivo, by measuring cFos activation; although cFos expression is not a linear readout of firing rates, the fraction of cFos-positive neurons in V1 reliably tracks overall levels of activity (Torrado Pacheco et al., 2019; Trojanowski et al., 2021; Wen et al., 2025; Wen and Turrigiano, 2021). To this end, we used the same post-learning treatment paradigm as above, but on day 3 allowed Sham, Hunt, or Hunt + XPro rats to forage for immobilized crickets in the hunting arena to ensure animals in all conditions were measured after a similar period of active exploration. Ninety minutes after the foraging session animals were sacrificed, and V1b was fixed and immunolabeled against NeuN (a pyramidal neuron marker, Bottorff et al., 2024; Chattopadhyaya et al., 2004; Nahmani and Turrigiano, 2014) and cFos (Figure 7I). Quantification of the fraction of NeuN positive neurons in L2/3 that were cFos positive revealed that this fraction was higher in the Hunt than Sham condition, and Hunt + XPro was significantly reduced relative to Hunt (Figure 7J). Taken together, our data show that learning induces slow, TNFα-dependent plasticity at excitatory synapses onto L2/3 pyramidal neurons to increase net local excitatory drive, and that this plasticity contributes to the post-learning increase in activity of L2/3 Pyramidal neurons.

### Blocking TNFα-dependent signaling after learning degrades the retention of hunting skills

If the slow, learning-induced increase in excitatory drive in V1b is important for consolidation of learning or maintenance of skills, then preventing it with XPro should also degrade behavioral performance. Consistent with this, XPro-treated mice performed significantly worse on day 6 than control mice (Figure 6G, Bissen et al., 2025). In our juvenile rat paradigm learning is complete in a single day (Figure 1E) and so can be cleanly separated from retention, so we next designed a paradigm to test for retention of hunting skills. Rats were allowed to hunt as normal on day 1, were injected with XPro or saline after the last hunting session (when learning was complete), and hunting behavior was tested 3 days later during two “probe” hunting sessions (probe, Figure 8A). As expected, both cohorts of animals performed similarly during learning (Figure 8C); however, the XPro cohort performed significantly worse during the probe sessions (Figures 8B, D-F). XPro-treated rats had roughly double the time to capture (Figure 8D), latency to attack (Figure 8E) and pursuit duration (Figure 8F), as control animals. Thus, loss of TNFα-dependent plasticity impairs the retention of hunting skills.

**Fig 8.**
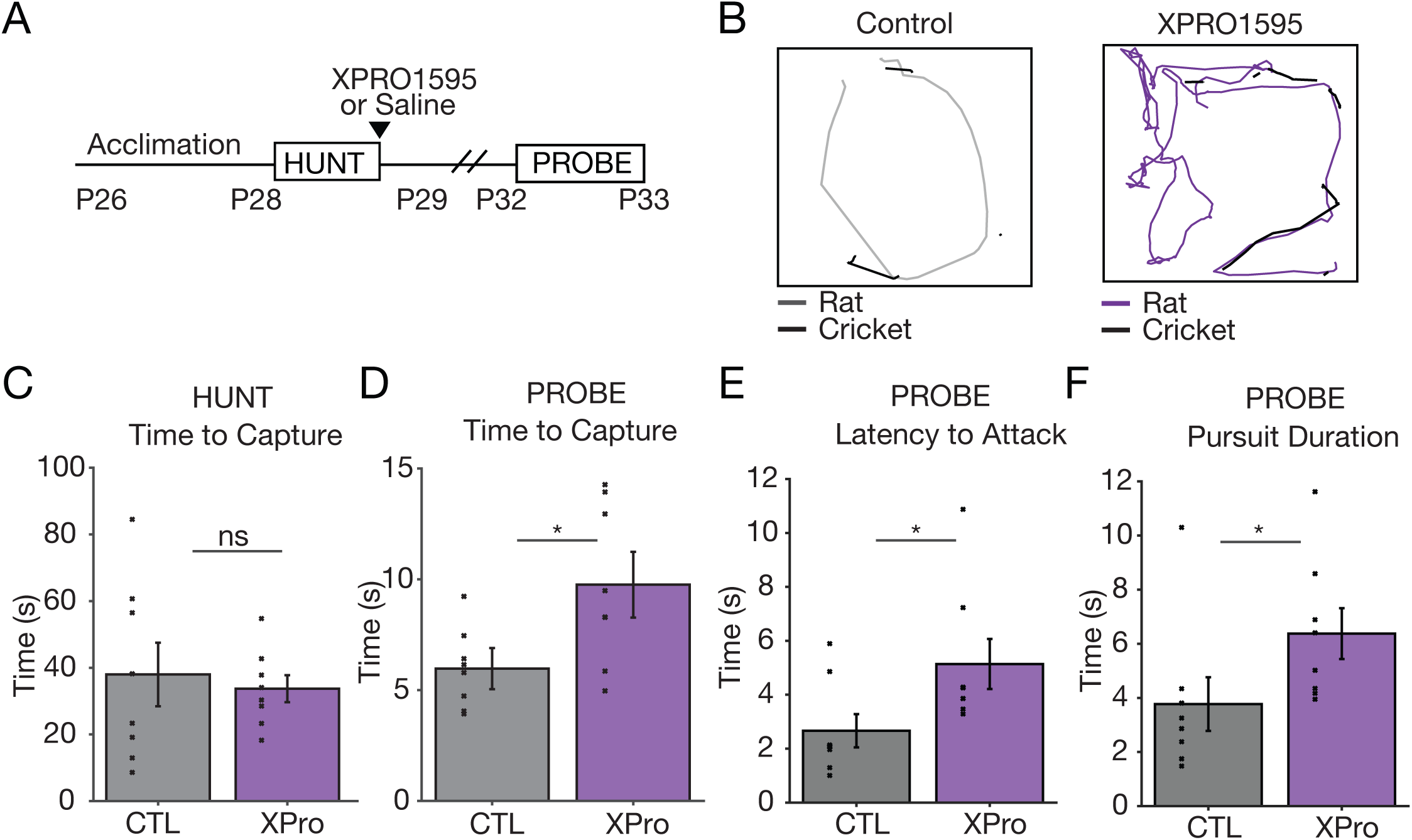
Retention of hunting skill is impaired by post-learning inhibition of TNFα-signaling. **A.** Experimental paradigm for determining the impact of XPRO1595 on retention of hunting skills; XPRO1595 was administered 1 hour following the completion of learning on Day 1, and behavior was subsequently measured in a probe session 3 days later. **B.** Left: Representative trajectory of control rat and cricket positions during probe sessions (grey: rat position, black: cricket position). Right: Representative trajectory of XPRO1595-injected rat and cricket positions during probe session (purple: rat position, black: cricket position). **C.** Average time to capture (s) on Day 1 for control (grey) and XPRO1595-injected (purple) rats. Individual points here and onward indicate animal averages. Here and below, CTL: n=8, XPro: n=8. Wilcoxon Rank Sum, p=0.7. **D-F**. (**D**) Average time to capture, (**E**) latency to attack, and (**F**) pursuit duration on Probe Day. Wilcoxon Rank Sum, time to capture, p=0.0004; latency to attack, p=0.04; pursuit duration, p=0.01.

## DISCUSSION

The idea that cortical neurons and circuits express stable activity set points to which they faithfully return when perturbed has become well-established (Hengen et al., 2016, 2013; Torrado Pacheco et al., 2021; Barnes et al., 2015; Wu et al., 2020; Dhawale et al., 2017; Jensen et al., 2022; Ma et al., 2019; Slomowitz et al., 2015; Ruggiero et al., 2025). In neocortex these set points are evident early in postnatal development and are stable through adulthood (Dhawale et al., 2017; Jensen et al., 2022; McGregor et al., 2024), suggesting they are stable features of neurons and/or cortical circuits. Recent work has shown that activity setpoints of hippocampal neurons can be altered through pharmacological manipulations (Slomowitz et al., 2015; Ruggiero et al., 2025; Styr et al., 2019) but whether these setpoints can serve as a substrate for plasticity during naturalistic learning was unknown. Here, we show that vision-dependent learning in juvenile rats induces a persistent resetting of visual cortical firing rates. First, we show rapid prey capture learning requires V1 and enhances the tuning of V1b neurons to specific hunting epochs. Chronic recordings before, during, and after this learning revealed a slow increase in baseline V1b firing rates that began shortly after learning was complete and persisted for days afterwards. This slow increase in firing was gated by sleep/wake states, and in L2/3 pyramidal neurons was driven by an increase in spine density and excitatory synaptic charge, with no change in net inhibition. Finally, inhibiting TNFα signaling post-learning prevented the enhancement of excitatory drive and reduced the retention of hunting skills. The slow timescale and dependence on behavioral state (Bottorff et al., 2024; Hengen et al., 2016; Torrado Pacheco et al., 2021), as well as reliance on TNFα signaling (Barnes et al., 2022, 2017; Kaneko et al., 2008; Pozo and Goda, 2010; Stellwagen and Malenka, 2006; Turrigiano, 2012; Wen et al., 2025), implicate homeostatic forms of plasticity in driving these changes. These results suggest that naturalistic learning in young animals can co-opt homeostatic forms of plasticity to drive neocortical networks to new activity setpoints.

Prey capture in rodents is a complex behavior that requires cooperation between many brain regions and circuits (Hoy et al., 2016, 2019; Zhao et al., 2019; Galvin et al., 2021; Johnson et al., 2021). Here, we focus on V1b because binocular vision is the primary sense used to track and grasp prey (Hoy et al., 2016, 2019; Michaiel et al., 2020; Galvin et al., 2021; Johnson et al., 2021; Holmgren et al., 2021), and we wished to understand whether vision-dependent learning during the visual system critical period can drive long-lasting plasticity within V1b that improves behavioral outcomes. We find that transiently inactivating V1 on the day of hunting slows initial time to capture and prevents the improvement in behavior across sessions; animals still orient towards and vigorously chase prey but are inefficient at tracking and capturing them (Movie S4). Multiple visual pathways likely cooperate to enable effective hunting; for example, the superior colliculus is also important for the execution of hunting skills (Hoy et al., 2019) and receives direct projections from V1 L5 (Hattox and Nelson, 2007; Kim et al., 2015; Liang et al., 2015), while providing feedback to visual cortical areas (Brenner et al., 2023). An interesting possibility is thus that prey capture learning enhances cooperation between cortical and subcortical visual circuits to effectively process behaviorally relevant visual stimuli.

In V1b, we find that neural responses become strongly modulated by pursuit across a single day of hunting, demonstrating rapid functional plasticity as performance improves. While movement speed does modulate V1b firing in these freely behaving juvenile rats, the magnitude of this effect is modest compared to head-fixed mice (Dipoppa et al., 2018; Liska et al., 2024; Niell and Stryker, 2010; Vinck et al., 2015), and saturates well below the average pursuit speed; thus simple changes in pursuit speed as animals learn cannot account for this plasticity. Interestingly, in critical period mouse V1, prey capture learning sharpens temporal frequency tuning in L2/3 but has little effect on other receptive field properties such as orientation tuning or binocular matching (Bissen et al., 2025), suggesting that the initial, rapid phase of V1 plasticity we observe here is likely tied to enhanced tracking of moving prey, rather than a general sharpening of classic receptive field properties. Recent work has shown that the firing of V1 neurons in freely behaving rodents cannot be well-explained by visual input alone, but instead is highly multimodal and likely represents active sensing that relies on integration of sensory and motor information to shape visual processing (Xu et al., 2023; Skyberg and Niell, 2024). In this context the development and amplification of reliable V1b responses during pursuit epochs as animals learn to hunt could reflect improved visuomotor processing for detection and tracking of prey (Guitchounts et al., 2020; Skyberg and Niell, 2024; Akitake et al., 2023; Histed, 2025).

Surprisingly, hunting induced the most dramatic changes in V1b activity after learning was complete. Despite animals returning to a home-cage environment with no further enrichment, both putative excitatory and inhibitory neurons underwent a slow increase in firing that persisted for at least 64 hours post-learning. This contrasts with chronic recordings from primary motor cortex in adult rats, where firing rates remained stable during and after motor learning (Dhawale et al., 2017; Jensen et al., 2022), and from the impact of passive sensory manipulations such as lid suture and eye-reopening in critical period V1, where firing rates (and other activity measures) are restored to baseline within a few days (Hengen et al., 2016, 2013; Keck et al., 2013; Torrado Pacheco et al., 2021; Barnes et al., 2015; Mrsic-Flogel et al., 2007; Bottorff et al., 2024). Similar to the impact of passive sensory manipulations, we find that sham hunting induced only a transient change in firing before activity returned to baseline. This suggests that naturalistic learning during the critical period has a unique effect on neocortex that drives a slow movement of network activity away from the default set point to a new, higher activity regime. This plasticity was especially evident in slowly-firing neurons, so it may serve to move a population of low information-containing neurons into a firing regime where they can contribute more effectively to sensorimotor processing and integration (Mizuseki and Buzsáki, 2013). Because our chronic recordings in these young, growing animals are limited in duration, we are unable to say whether this new activity state is permanent or merely long-lasting.

Homeostatic plasticity is generally understood to stabilize activity around an established activity setpoint (Turrigiano and Nelson, 2004; Abbott and Nelson, 2000; Turrigiano, 2012; Keck et al., 2017); in contrast, here we find that after learning, network activity slowly moves away from the original setpoint to a new, higher activity state. If this slow plasticity arises through a rapid, learning-induced resetting of homeostatic setpoints, and then a slow movement to these new setpoints, then this process should unfold according to the rules and mechanisms of homeostatic plasticity. Plasticity of excitatory synapses is a major contributor to firing rate homeostasis within V1 (Keck et al., 2013; Lambo and Turrigiano, 2013; Torrado Pacheco et al., 2021; Turrigiano, 2012; Wen and Turrigiano, 2021, 2021; Wu et al., 2020), and expression of upward forms of excitatory homeostatic plasticity (Barnes et al., 2022, 2017; Stellwagen and Malenka, 2006; Steinmetz and Turrigiano, 2010; Wen et al., 2025; Kleidonas et al., 2023), as well as the restoration of network activity (Barnes et al., 2022, 2017; Kaneko et al., 2008) depend on TNFα signaling. Synaptic homeostasis can target both strength and number of synapses (Turrigiano et al., 1998; Wierenga et al., 2006; Keck et al., 2017; Desai et al., 2002; Guerrero and Turrigiano, 2025; Lu et al., 2025), and we find the density of excitatory synapses onto L2/3 pyramidal neurons increases with network activity in a TNFα-dependent manner. As other forms of cortical synaptic potentiation, such as LTP, are insensitive to TNFα blockade (Kaneko et al., 2008; Stellwagen and Malenka, 2006), this strongly implicates homeostatic, rather than Hebbian, synaptic plasticity. On the other hand, unlike for deprivation-induced homeostatic regulation (Wen and Turrigiano, 2021; Lambo and Turrigiano, 2013), intrinsic excitability of L2/3 neurons was not enhanced post-learning. These differences are consistent with previous work showing that which cellular homeostatic mechanisms are recruited depends on many factors, including when and how the network is perturbed (Desai et al., 2002; Maffei and Turrigiano, 2008; Wen et al., 2025). In sum, the synaptic changes set in motion in L2/3 after naturalistic learning are slow, widespread across the dendritic arbor (Figure 7D), result in a net increase in excitatory drive, and are TNFα-dependent, all defining features of homeostatic synaptic plasticity.

The homeostatic nature of the post-learning firing rate increase is further supported by its behavioral state-dependence; like upward firing rate homeostasis induced by visual deprivation (Hengen et al., 2016; Bottorff et al., 2024), the post-learning increase in firing accumulated incrementally across successive periods of wake. This contrasts with Hebbian potentiation in V1, which is either independent of sleep/wake states or occurs selectively during sleep(Aton et al., 2014, 2009; Dumoulin Bridi et al., 2015; Frank, 2017). Interestingly, memory consolidation in the hippocampus (Aton et al., 2014; Diba and Buzsáki, 2007; Frank et al., 2001; Ji and Wilson, 2007; Josselyn and Tonegawa, 2020; Lee and Wilson, 2002; Wilson and McNaughton, 1994) (but see: Carr et al., 2011; Jadhav et al., 2012; Tang et al., 2017; Shin et al., 2019) and some forms of experience-dependent plasticity consolidation in V1 (Aton et al., 2014; Frank, 2017) occur preferentially during sleep states, whereas here we find that consolidation and retention is associated with net upward wake-dependent plasticity. Although the net effect on firing over time was upward, there was a pronounced oscillation in firing that was associated with cycles of wake (upward changes in firing rate) and sleep (downward changes in firing rate), raising the possibility that wake-dependent and sleep-dependent mechanisms cooperate after learning to reshape network activity as hunting skills are consolidated.

What function(s) might this resetting of network activity serve? In insular cortex, homeostatic plasticity can enable the emergence of memory specificity after gustatory learning (Wu et al., 2021), suggesting that task-specific processing can be maintained or even enhanced during resetting of network activity. By bringing very low firing rate neurons into a more functional firing rate regime, this resetting may better integrate these neurons into the local network to enhance sensory processing during complex behaviors and thus contribute to the consolidation of the visuo-motor skills needed for successful hunting. Another interesting possibility is that these newly active neurons might increase the pool of neurons available as substrates for learning, and thus provide a future advantage when animals attempt to learn new vision-dependent skills. Most broadly, this work shows that neocortical activity setpoints are malleable and suggests that learning-dependent changes in these setpoints contributes to behavioral improvements during naturalistic learning.

## STAR Methods

### KEY RESOURCES TABLE

| REAGENT OR RESOURCE | SOURCE | IDENTIFIER |
| --- | --- | --- |
| **Antibodies** |  |  |
| Rat monoclonal anti-mCherry | ThermoFisher | RRID: AB_2536611 |
| Mouse monoclonal anti-NeuN | Millipore Sigma | RRID: AB_2736207 |
| Rabbit monoclonal anti-cFos | Cell Signaling Technologies | RRID: AB_2298772 |
| Goat polyclonal anti-mouse Alexa 488 | ThermoFisher | RRID: AB_2534069 |
| Goat polyclonal anti-rat Alexa 594 | ThermoFisher | RRID: AB_10561522 |
| Goat polyclonal anti-rabbit Alexa 647 | ThermoFisher | RRID: AB_2535813 |
| Chicken polyclonal anti-GFP | Millipore | RRID: AB_90890 |
| Rabbit polyclonal anti-NeuN | ThermoFisher | RRID: AB_2736207 |
| Goat polyclonal anti-chicken Alexa 488 | ThermoFisher | RRID: AB_2534096 |
| Goat polyclonal anti-rabbit Alexa 568 | ThermoFisher | RRID: AB_143157 |
| **Bacterial and virus strains** |  |  |
| AAV9.hSyn.DIO.hM3D(Gq).mCherry | Addgene | RRID: Addgene_44361 |
| **Chemicals, peptides and recombinant proteins** |  |  |
| XPro1595 | INmune Bio | N/A |
| Clozapine-N-Oxide (CNO) | HelloBio | N/A |
| **Experimental models: Organisms/strains** |  |  |
| Mouse: B6.Cg-Tg(Thy1-YFP)HJrs/J | Feng et al, 2000 | RRID: IMSR_JAX:003782 |
| Rat: PvalbCre | Yu et al., 2018 | N/A |
| Rat: Long Evans | Charles River Laboratories | RRID:RGD_60991 |
| **Software and algorithms** |  |  |
| Synapse | Tucker Davis Technologies | RRID:SCR_006495 |
| DeepLabCut | Mathis et al., 2018; Nath et al., 2019 | RRID:SCR_021391 |
| Huygens Essential | Scientific Volume Imaging | RRID: SCR_014237 |
| ImageJ/Fiji | Fiji | RRID: SCR_002285 |
| Imaris | Oxford Instruments | RRID: SCR_007370 |
| GraphPad | Prism | RRID: SCR_002798 |
| Matlab | MathWorks | RRID: SCR_001622 |
| Illustrator | Adobe | RRID: SCR_010279 |
| Photoshop | Adobe | RRID: SCR_014199 |
| Custom MATLAB software | This manuscript | Lab GitHub |

### EXPERIMENTAL MODEL AND SUBJECT DETAILS

#### Animals

All procedures were approved by the Brandeis University Institutional Animal Care and Use Committee and conformed to the National Institutes of Health *Guide for the Care and Use of Laboratory Animals*; colonies of rats and mice were maintained at Brandeis in the Foster Biomedical Research Facility. For all experiments, rats and mice of both sexes were used. Rats were juvenile Long Evans (Charles River Laboratories, Cambridge, MA, USA) or Parvalbumin (Pvalb)-Cre on a Long Evans background (Yu et al., 2018), founder rats obtained from Loren Frank (University of California, San Francisco). Mice were Thy1-YFP (B6.Cg-Tg(Thy1-YFP)HJrs/J, The Jackson Laboratory, Bar Harbor, ME, USA), which express the yellow fluorescent protein (YFP) in a subset of V1b L2/3 pyramidal neurons.

### METHOD DETAILS

#### Rat prey capture learning paradigm

Littermate rats were transferred to behavioral testing rooms 2 days prior to cricket hunting. Rats were given standard housing, chow, and nesting materials as previously described (Bottorff et al., 2024; Hengen et al., 2016, 2013; Torrado Pacheco et al., 2021). A circadian light cycle of 12-hour light/dark periods was maintained, with dark hours between 7:30am and 7:30pm. All cricket hunting experiments took place in a behavioral testing arena as described previously (Groves Kuhnle et al., 2022; Bissen et al., 2025). Briefly, the behavioral arena consisted of a 14”x14”x14” plexiglass chamber with a removable floor with absorbent pad, social partner divider, and detachable cricket dispensers. In “home cage” configuration, rats were given *ad libitum* chow, water, bedding, and access to a social partner via a perforated, transparent divider (Figure 1B. In “hunting configuration,” the social partner was removed, chow, water, and nesting materials were removed, the floor was replaced with a clean pad, and cricket dispensers were affixed to the four corners of the arena (Figure 1A). Additionally, diagonal stripes were attached to the outside of the arena to provide visual distractors on all four sides of the testing arena. For two days, animals were handled by experimenters and acclimated to the behavioral arena in “hunting configuration” for 30 minutes and fed two live, but immobilized crickets each day. Crickets were sourced from Fluker Farms (Port Allen, LA, USA). On the second day of acclimation, immediately prior to lights out, rats were food deprived for no more than 16 hours. Behavioral testing began the next day, when the behavioral arena was configured into “hunting” mode and crickets were loaded into the cricket dispensers. Each hunting session consisted of a 5-minute baseline period prior to when the first cricket was dispensed. Crickets were dispensed using custom, in-house designed 3D printed dispensers controlled by custom Arduino-powered stepper motors and software. With each cricket hunt, dispensers on opposite sides of the arena were programmed to turn to prevent the animals from predicting which dispenser a cricket would be released from. Behavior was recorded at 20 frames per second using either a Logitech C922x HD Pro Webcam or an infrared-capable Sony IMX323 webcam, via Synapse (Tucker Davis Technologies (TDT), Alachua, FL, USA) software. Each hunting session consisted of 10 crickets and typically lasted 30-45 minutes. After each hunting session, the arena was returned to “home cage” configuration. For all hunting experiments, rats performed 3 hunting sessions on Day 1. In a subset of experiments, animals performed a probe day of hunting 3 days later (Day 4), consisting of prior food deprivation and 2 hunting sessions. At the completion of the final hunting session of the day, animals were returned to *ad libitum* food and water access. On days where hunting did not occur, animals were housed in “home cage configuration.” Animals were sacrificed following the end of the behavioral paradigm in accordance with Brandeis Institutional Animal Care and Use Committee guidelines.

#### Mouse prey capture learning experiments

For mouse prey capture experiments, we used the paradigm described in Bissen et al., (Bissen et al., 2025), modified from Groves Kuhnle et al., (Groves Kuhnle et al., 2022), with the same arena configuration as for rats. Briefly, mice were transferred to behavioral testing rooms 2 days prior to the first day of hunting. Mice were handled and habituated for 1 hour and fed one live, but immobilized cricket per day. Mice were food deprived for up to 16 hours prior to the start of each hunting/sham session but had water ad libitum. Mice then underwent three days of one session per day of hunting/sham (days 1-3), followed by two days with no hunting/sham session (days 4-5) and one last day with one hunting/sham session (day 6). Each hunting session comprised 10 live crickets, while each Sham session comprised 10 immobilized crickets. All hunting/sham sessions took place over the course of 6 days between P28 and P35, with the first session taking place on P28-P30.

#### Chemogenetic (DREADDs) disruptions of Visual Cortex (V1)

Virus injections of excitatory DREADDs (AAV9-hSyn-DIO-hM3D(Gq)-mCherry, Plasmid #44361-AAV9, Addgene, Watertown, MA, USA) were performed on juvenile PV-Cre^+/-^and PV-Cre^-/-^ littermates ages P14-P18(Yu et al., 2018). Rats were anesthetized with vaporized isoflurane (4-5% induction, 1-2% maintenance) via an integrated vaporizer (Somnosuite, Kent Scientific, Torrington, CT USA) connected to a stereotaxic holder (Model 923-B with Model 1924-C-11.5 mask, Kopf Instruments, Tujunga, CA. Two injection sites were chosen per hemisphere, and 200-300nL of virus was delivered into V1 via a micropipette and oil-hydraulic injector (MO-10, Narishige International USA, Amityville, NY, USA). Clozapine N-Oxide (CNO, HelloBio, Princeton, NJ, USA) was dissolved in water and 10mM saccharine chloride was added to reach a final concentration of 0.05mg/mL (Bottorff et al., 2024). At lights out on the evening prior to hunting, rats’ drinking water was replaced with CNO-water, and access was given until the cessation of hunting session 3 the following day. Animals drank 15-20 mL over the course of administration to reach a desired CNO dose of 10 to 15 mg/kg (Bottorff et al., 2024).

#### Continuous-single cell recordings in freely behaving animals

Juvenile Long Evans rats (P22-P24) were implanted with electrode arrays as previously described (Hengen et al., 2016; Torrado Pacheco et al., 2019, 2021; Bottorff et al., 2024). In brief, custom 16 or 32-channel microwire arrays (tungsten, 33-micron diameter, Tucker Davis Technologies (TDT), Alachua, FL, USA) were implanted in either V1b or V1m, with location validated post-hoc through histological reconstruction. Rats were anesthetized with vaporized isoflurane (4-5% induction, 1-2% maintenance) via an integrated vaporizer (Somnosuite, Kent Scientific, Torrington, CA, USA) connected to a stereotaxic holder (Model 923-B with Model 1924-C-11.5 mask, Kopf Instruments, Tujunga, CA, USA). The skull was exposed via a small incision, cleaned with hydrogen peroxide, and bleeding spots were cauterized. Bone screws (Antrin Inc, Fallbrook, CA, USA) were implanted above the cerebellum posterior to bregma and bilaterally above motor cortex. V1b or V1m were targeted using stereotaxic coordinates and anatomical landmarks, and the bone was removed using a 27G needle. A headcap was created using UV-cured acrylic (Flow-It Flowable Composite, Pentron Technologies, Orange, CA, USA). The dura was removed using a 27G syringe and kept moist using chilled, bacteriostatic sterile 0.9% saline (Hospira Ince, Lake Forest, IL, USA). Electrodes were slowly lowered into the brain, and the craniotomy was encapsulated with a silicone elastomer (Kwik-Cast, World Precision Instruments, Sarasota, FL, USA). The electrode array was secured using additional UV-cured acrylic. A steel ground wire from the electrode array was then secured to one of the bone screws over motor cortex using solder paste and additional dental acrylic. Finally, a braided steel wire from the electrode array was implanted into the nuchal muscle for EMG recordings.

Following electrode implantation, animals were allowed to recover for at least 2 days with *ad libitum* access to food and water. During this time, animals were administered postoperative injections of meloxicam and penicillin during the first two days of recovery and were handled daily by experimenters. Animals were then transferred to our custom, convertible behavioral arena in the previously “home cage” configuration and allowed to habituate to the arena for two days prior to initiation of hunting, as described above. The microwire arrays were connected to TDT commutators via ZIF-clip headstages that enable animals to move freely throughout the arena. Implanted animals were accompanied by a littermate social partner in the adjacent chamber in “home configuration.” Animals were continuously connected via an active commutator for the duration of the experiment and behavior was continuously recorded using an IR-capable camera, enabling constant recording of electrophysiology and behavioral data. Electrophysiology data was acquired at 25kHz and streamed using a TDT Neurophysiology Workstation and Data streaming pipeline. Spike extraction, clustering, and sorting were performed using custom MATLAB and Python code as previously described (Hengen et al., 2016; Torrado Pacheco et al., 2019, 2021; Bottorff et al., 2024).

#### Slice electrophysiology recordings

Rats were deeply anesthetized with isoflurane and rapidly decapitated. Brains were rapidly removed and sliced in ice cold, oxygenated artificial cerebrospinal fluid, (aCSF, in mM: 126 NaCl, 25 NaHCO3, 3 KCl, 2 CaCl2, 2 MgSO4, 1 NaH2PO4, 0.5 Na-Ascorbate, osmolarity adjusted to 315 mOsm with dextrose, pH 7.35) using a vibratome (Leica VT1200S). Slices were allowed to recover in oxygenated aCSF at 34°C for 1 hour prior to the start of recordings, then moved to room temperature. All recordings were performed within 6 hours of slicing. During recordings, slices were continuously perfused with oxygenated aCSF at 34°C. For intrinsic excitability recordings, aCSF contained synaptic blockers (in mM: 25 DNQX, 50 APV, 25 PTX) and used K-gluconate internal (in mM: 100 K-gluconate, 20 KCl, 10 HEPES, 5.37 Biocytin, 10 Na-Phosphocreatine, 4 Mg-ATP, and 0.3 Na-GTP, with sucrose added to bring osmolarity to 295 mOsm and KOH added to bring pH to 7.35). For spontaneous mEPSC recordings, aCSF contained synaptic blockers (APV and PTX) at the same concentrations above as well as TTX to block action potentials (0.2 uM), with a CS based internal solution (in mM: 115 Cs-Methanesulfonate, 10 HEPES, 10 BAPTA•4Cs, 5.37 Biocytin, 2 QX-314 Cl, 1.5 MgCl2, 1 EGTA, 10 Na2 -Phosphocreatine, 4 ATP-Mg, and 0.3 GTP-Na, with sucrose added to bring osmolarity to 295 mOsm, and CsOH added to bring pH to 7.35). Slices were visualized on an Olympus BX51WI upright epifluorescence microscope using a 10x air (0.13 numerical aperture) and 40x water-immersion objective (0.8 numerical aperture) using infrared differential interference contrast optics and a CCD camera. Pyramidal neurons in L2/3 were targeted visually and confirmed with post-hoc reconstruction using biocytin labeling. Glass pipettes were pulled using a Sutter P-97 to obtain a tip resistance between 3-6 MOhm. Data was acquired at 10 kHz using Multiclamp 700B amplifiers and CV-7B headstages (Molecular Devices, Sunnyvale CA), with MATLAB-based data acquisition software, WaveSurfer (v 1.0.5, Janelia, Ashburn VA). All post-hoc analysis was performed using custom scripts written in MATLAB (Mathworks, Natick MA).

For spontaneous synaptic current recordings, slices were incubated in active ACSF (in mM: 126 NaCl, 25 NaHCO_3_, 3.5 KCl, 0.5 MgCl_2_, 1 CaCl_2_, 1 NaH_2_PO_4_, 0.5 Na-Ascorbate, osmolarity adjusted to 315 mOsm with dextrose, pH 7.3) in the absence of any pharmacological agents, and the Cs-based internal recording solution was used. For each neuron, the inhibitory reversal potential was determined by first holding the cell at -55 mV, then decreasing the holding potential in 3-5 mV increments until outward currents were no longer detectable. Excitatory synaptic currents were then recorded by holding the neuron at the experimentally determined inhibitory reversal potential (between -55 mV and -70 mV). The excitatory reversal potential was determined in the same manner, starting at -10 mV, and inhibitory synaptic currents were recorded at the experimentally determined excitatory reversal potential (between -5 mV and +10 mV). Recordings were taken in 30 second traces so that passive properties could be continuously monitored; total recording length for each neuron was 3-5 minutes for both excitatory and inhibitory currents.

#### Immunohistochemistry

##### Rat Immunohistochemistry

Animals were deeply anesthetized with a Ketamine/Xylazine/Acepromazine (KXA) cocktail (70 mg/kg ketamine; 3.5 mg/kg xylazine; 0.7 mg/kg acepromazine) and perfused with phosphate-buffered saline (PBS) and 4% paraformaldehyde. Fixed brains were sliced at 60-micron thickness on a vibratome (Leica VT1200, Leica Biosystems, Deer Park, IL, USA) and stored in 1x PBS. For electrophysiology experiments, brains were stained with DAPI and imaged with an epifluorescence widefield microscope (Keyence BZ-X, Keyence, Woburn, MA, USA) to verify electrode location post-hoc. For DREADDs-manipulation experiments, slices were washed, incubated in a blocking solution for 2 hours, then incubated with primary antibodies: rabbit anti-cFos (9F6, Cell Signaling Technology) (1:200), rat anti-mCherry (M-11217, Invitrogen) (1:1000), and mouse anti-NeuN (MAB-377, Millipore Sigma) (1:500). The following day, slices were incubated with secondary antibodies: goat anti-mouse Alexa-488 (A-11001, Invitrogen), goat anti-rat Alexa-594 (A-11007, Invitrogen), and goat anti-rabbit Alexa-647 (A-21245, Invitrogen) (all 1:400). Images were obtained using a confocal microscope (Leica SP5, Leica Microsystems, Deer Park, IL, USA).

##### Mouse Immunohistochemistry

Mice were sacrificed a minimum 3 hours after the last hunting session on day 6 as for rats (described above). After dissection and post-fixation, brains were sectioned in 200µm thick slices to encompass complete dendritic trees. Selected slices containing entire dendritic arbors of L2/3 pyramidal neurons in V1b were permeabilized, blocked, and incubated with primary antibodies in blocking solution for 48-72 hours at 4 °C, and incubated with corresponding secondary antibodies for 2 hours at room temperature. Slices were stained with chicken anti-GFP (90890, MilliporeSigma) (1:500) and rabbit anti-NeuN (2736207, ThermoFisher) (1:500), followed by goat-anti-chicken Alexa 488 (143165, ThermoFisher) and goat-anti-rabbit Alexa 568 (143157, ThermoFisher) (both 1:500). Images were obtained using a confocal microscope (LSM 880, Zeiss, Oberkochen, Germany).

#### Pharmacological Manipulations using XPRO1595

The pegylated protein tumor necrosis factor alpha (TNFα)-1 inhibitor XPro1595 (Zalevsky et al., 2007) (INmuneBio, Boca Raton, FL, USA) was administered as previously described (Bissen et al., 2025; Barnes et al., 2022, 2017; Wen et al., 2025; Barnum et al., 2014). XPRO1595 was dissolved in 0.9% saline to the desired concentration (1mg/mL). Animals were weighed and the appropriate amount of XPRO1595 was delivered via subcutaneous injection to reach the desired dosage (10mg/kg). For mouse cricket hunting experiments, XPRO1595 was administered twice, once approximately one hour following the Day 1 hunting session, and again 3 days later, roughly halfway through the paradigm, after the Day 3 hunting session. The control cohort consisted of non-injected (n=5) and saline-injected (n=3) mice; these two cohorts had similar learning curves (see Bissen et al., 2025, where L5 pyramidal neurons from the same animals were analyzed) and were therefore combined. For rat cricket hunting experiments XPRO1595 was administered to rats once, approximately one hour following the cessation of hunting session 3 on the first day of hunting. Animals were retested 3 days later on “Probe” day, consisting of 2 hunting sessions of 10 crickets/session.

### QUANTIFICATION AND STATISTICAL ANALYSIS

#### Prey Capture Learning Behavioral Scoring

##### Quantification of Hunting Performance

Behavior was recorded during each cricket hunting session were manually scored to quantify cricket entrance, the pursuit initiation, cricket capture, and end of consumption. Time to capture was calculated as the time (in seconds) from cricket entrance to capture. Latency to attack was calculated as the time from cricket entrance to pursuit initiation, and attack duration was calculated from pursuit initiation to capture. Further behavioral analysis was performed using DeepLabCut (DLC,Mathis et al., 2018; Nath et al., 2019) to assess differences between conditions in DREADDs-manipulation experiments, as well as in XPRO1595 experiments. Over 300 still frames from hunting videos were chosen and manually labeled with cricket position and the position of various rat body features. As previously described (Groves Kuhnle et al., 2022; Bissen et al., 2025), a ResNet50 neural network was trained on these manually labeled datasets. Following training, the performance of the network was evaluated using test data. Outlier frames were then chosen and manually labeled for subsequent training iterations to improve performance. DeepLabCut outputs of cricket position coordinates and rat head position were then used to create trajectories of cricket hunting pursuits using custom software (MATLAB).

##### Sleep-Wake Behavioral State Scoring

Behavioral state scoring was accomplished as previously described using a semi-automated MATLAB GUI (Cary and Turrigiano, 2021; Torrado Pacheco et al., 2021) with the modification that animal movement was tracked using DLC (Mathis et al., 2018; Nath et al., 2019). Briefly, local field potentials were extracted from 3 of the highest quality channels, downsampled, and averaged. We computed the spectrograms in 5 second bins from 0.3 to 15 Hz. Standard delta (0.3 – 4 Hz) and theta (5 – 8 Hz) bands were used to differentiate cortical states. Animal movement was measured using downsampled mean EMG signal and DLC tracked body poses separate for the head and body of the animal. DLC poses were processed by first removing all low confidence points (<0.9 confidence). 8 head features were tracked (nose, ears, etc.) along with 3 body features, allowing for separate head/body movement tracking. The instantaneous speed of each pose was calculated and smoothed with a 30 frame lowess function (MATLAB). The head and body speed were determined by taking the median speed of each pose. A trained human scorer used the GUI to manually score the first 10 hours of behavioral data, and then subsequently supervised the labeling of the rest of the experimental data with a random forest model trained on those first 10 hours (process was repeated for each animal).

##### Spike Extraction, Clustering, and Spike Sorting

Spike extraction, clustering, and sorting were done as previously described (Hengen et al., 2016; Torrado Pacheco et al., 2021; Bottorff et al., 2024; Torrado Pacheco et al., 2019). Briefly, putative spikes were identified by greater than or equal to 4 standard deviations from mean signal. Using principal component analysis (PCA), the spikes from each channel were clustered using KlustaKwik (Harris et al., 2000).Clusters were classified and scored using a random forest model based on 1200 manually scored clusters encompassing 19 features (e.g. interspike interval (ISI) contamination, waveform template matching, waveform kinetics, and noise contamination). Only single-unit clusters that were classified as high quality were used for our firing rate analyses. Clusters were further classified by their putative cell type, regular spiking (RSU) or putative fast spiking (FSU), using established criteria (Hengen et al., 2016; Torrado Pacheco et al., 2021; Bottorff et al., 2024; Torrado Pacheco et al., 2019).

##### Hunting Firing Rate Modulation

To determine which hunting behaviors significantly modulate firing rates, we took a bootstrap reshuffling approach. Firing rates were smoothed with a Gaussian kernel (σ = 3s for RSU and 5s for FSU cells; chosen empirically based on FR variance during this behavior while still capturing true FR dynamics). The behavior times for a whole hunting session (10 crickets) were shifted randomly (retaining the relative behavior time intervals) within an expanded time window (+1 hour on either side). This shuffling was done 500 times for each cell. For each shuffle, the FR_ICI_, FR_pursuit_, FR_consumption_, was determined for every cricket. Both FR_pursuit_ and FR_consumption_ were normalized to the corresponding prior FR_ICI_. This created a shuffle control distribution, where the actual normalized FR_pursuit_ and FR_consumption_ when above 95^th^ or below 5^th^ percentile were deemed significant (upward and downward, respectively). Firing rate histograms around pursuit start and cricket capture were made by aligning a 30 second window around those times and normalizing to an 8 second baseline period at the beginning. This was done to focus on the FR dynamics specifically around those short behavioral moments. The FR plots with normalized pursuit time were created by linearly interpolating 1000 points (bin size = 0.1 seconds) for each pursuit regardless of duration, normalizing to an 8 second baseline period before pursuit, and applying moving average filter (window = 50).

##### Hunting Movement Metrics

Animal position/movement from DLC was processed as described above. Average pursuit speeds were calculated over the entire pursuit interval for each cricket/animal. Max speeds were taken as the instantaneous smoothed speed across interval. Distance traveled was calculated as integral of the speed (0.05 second bin). The FR versus speed plot was created by comparing speeds and FRs (spike time histograms also with 0.05 second bins, smoothed with Gaussian kernel σ = 3s), during the light period of the hunting day (the ∼4-hour hunting period with ∼4 hours preceding it). Average trace was computed from the cell averages.

##### Timepoint Analysis of Ensemble Firing Rates

To calculate firing rates at discrete timepoints, we clustered spiking data in 6-hour bins from each animal at the indicated timepoints. We then examined each putative cluster and identified high-quality clusters at each timepoint as described above. We calculated firing rates by taking the inverse of the mean interspike interval (ISI) for each cluster.

##### Continuous Single Unit Analysis

To follow single units over time, we spike-sorted on the entire continuous recording and then determined “on” and “off” times for each high-quality cluster to determine when well-isolated units could be detected, as previously described (Hengen et al., 2016; Torrado Pacheco et al., 2021; Bottorff et al., 2024). Briefly, we first examined each high-quality cluster’s stability by examining its hourly firing rate and ISI contamination, restricting “on” times to periods when ISIs < 2.5ms were below 4%. Next, we calculated the sum of squared errors (SSE) of the waveform by comparing each hour of the recording to the average waveform across the entire experiment, to ensure waveform shape and peak amplitude remained stable. We manually inspected the hourly SSE to carefully detect deviations from the average waveform and exclude these times from our analyses. Using this combination of metrics, we assigned “on” and “off” times when each cluster was well-isolated and stable, and considered only units that were well-isolated and had stable SSE for >70% of the experiment duration. To calculate each unit’s firing rates, we computed spike counts in 60s bins and applied a Gaussian kernel with σ = 300 s (see Torrado Pacheco et al., 2021). For timepoint measurements of continuously recorded cells (Figure 3D-H, Figure 4, we averaged continuous firing rates for each cluster across 6hr bins at each timepoint.

##### Behavioral State Analysis

To measure sleep/wake effects on firing rates (FRs) we first detected extended sleep/wake episodes. Episodes were defined as periods of near-continuous (<120 second interruptions) sleep/wake lasting at least 30 minutes. To determine overall FR change (Figure 5B, E), we measured the difference in FRs in flanking windows located in the first 30% and last 30% of each episode. We calculated z-score change by first normalizing 𝑧𝐹𝑅=(𝐹𝑅–𝜇)/𝜎, where 𝜇 (mean) and 𝜎 (standard deviation) were calculated based on the FR data only within the episode. Next, we found the mean change 𝐹𝑅_last_–𝐹𝑅_first_ where both were mean FRs for the same given state in the flanking periods (Figure 5B). Percent change was calculated %change=100*(𝐹𝑅_last_– 𝐹𝑅_first_)/ 𝐹𝑅_first_, where both 𝐹𝑅_last_ and 𝐹𝑅_first_ were mean FRs for the same given state in the flanking periods.

To determine how the behavioral state regulation of FRs unfolds across the post hunt period (Figure 5E), we divided the timeline into two 24-hour bins (centered at 12hr and 36hr) and a 12 hour (centered at 60hr) final period. FR change correlation plots (Figure 5C, D) were made by a similar method but instead mean FRs were found for each instance of a behavioral state (NREM, REM, quiet/active wake) across the extended period (no flanking window restriction). That is, wake percent change was calculated %change=100*(𝐹𝑅_n_– 𝐹𝑅_first_)/ 𝐹𝑅_n_, where 𝐹𝑅_n_ was the mean FR for an instance of either active/quiet wake in the period and 𝐹𝑅_first_ was the mean FR for the first quiet wake in that same period. Sleep percent change was calculated relative to the first NREM FR.

#### Confocal image acquisition and analysis

##### cFos Quantification

CFos expression in PV-Cre^+^ and PV-Cre^-^ animals was quantified as previously described (Wen and Turrigiano, 2021; Bottorff et al., 2024). We created maximum projection images from z-stacks taken with a 20x objective in V1 of fixed slices from both cohorts of animals. Images were background subtracted with a rolling ball radius of 50 pixels. NeuN neurons were traced manually from the NeuN channel, and the cFos+ neurons were identified by thresholding in the cFos channel of the image. NeuN labeled and NeuN/cFos double-labeled neurons were quantified from all V1 images to find a total ratio of (NeuN+cFos+ neurons/ total NeuN+ neurons) per animal using ImageJ (NIH, Washington, DC).

##### Dendritic Spine Quantification

High-resolution, large-scale imaging of spine density and morphology in L2/3 pyramidal neurons was performed using an inverted confocal laser scanning microscope (LSM880, Zeiss). The localization of pyramidal neurons to V1b was first confirmed with a 10x dry objective (NA 0.45)., and analysis was confined to neurons where the full dendritic arbor could be reconstructed. The entire apical and basal trees of each neuron were imaged using a 63x oil objective (NA 1.4) at 1024 x 1024 pixels (pixel size of 220 x 200 x 300nm (Day 6, Day 6 + XPro) or 90 x 90 x 300nm (Day 1)). To improve resolution and signal to noise ratio, images were deconvolved in Huygens (Scientific Volume Imaging) using the classic maximum likelihood estimation (CMLE) algorithm. The entire dendritic trees and all spines were traced manually using the Filament Tracer module in Imaris (Oxford Instruments). Dendritic spine density was calculated for each dendritic segment, and densities and spine head diameters were obtained using the Statistics module in Imaris.

Representative images were improved for clarity by removing signals from other dendrites in Imaris as follows: the selected dendrite was traced as a new filament and used to reconstruct a surface with a low intensity threshold using the Surface module. This artificially enlarges the surface so that it becomes a cylinder encompassing the dendritic shaft and all the attached spines. The fluorescence signal of all pixels outside of that surface were then set to 0 and in effect masked, so that only the dendritic stretch of interest would be shown in the representative picture. Brightness and contrast were subsequently enhanced using the unsharpen filter in Photoshop (Adobe).

##### Synaptic charge measurements

To accurately calculate synaptic charges, baseline currents were first carefully determined. Each trace was first corrected for baseline drift and then divided into 1 second segments, and a histogram was generated for all current values in this segment. The peak of the histogram was defined as the “corrected” baseline, and noise values should be symmetrically distributed on its two sides. For excitatory currents, all values on the positive side were considered as noise, and the total excitatory charge was calculated by taking the integral of values on the negative side and subtracting the integral of values on the positive side (putative noise). This was then repeated for all segments and all traces, and the charge values were averaged to generate the mean value for the cell. Inhibitory charges were determined similarly by regarding current values on the negative side as noise. To ensure baselines were stable throughout the recording, a histogram was generated for all baseline values from a given cell. Segments with baseline values whose z-scores were greater than 4 were excluded.

Well-isolated events were automatically detected using in-house scripts written with MATLAB as previously described (Wen et al., 2025). Briefly, the script slides an event-shaped template to identify putative events, which were then passed through multiple quality control modules to exclude false-positives and bursts. Mean amplitude and frequency were first calculated for each 30 s recording segment, which were then averaged to generate the mean value for each cell.

### Statistical Analyses

Data analysis was performed using Graphpad Prism or custom code written in MATLAB (see Key Resources Table, available on GitHub). Values were reported in the text body as mean ± standard error of the mean (SEM). Individual data points represent session averages on behavioral curves, individual animals on behavioral plots, and individual neurons on firing rate plots except where noted otherwise. Parametric or non-parametric tests were chosen depending on whether data passed normality tests. To compare the means of two non-normally distributed groups, we used a Wilcoxon Rank Sum test with Bonferroni correction. To compare pairs of non-normally distributed data, we used a Wilcoxon Sign Rank test with Bonferroni correction. To compare differences within a single group across multiple timepoints, we used a Friedmann’s test with Dunn correction; between two groups across multiple timepoints, we used a two-way ANOVA with Tukey-Kramer correction. Comparison of cumulative distributions was done with Kolgornov-Smirnov or Anderson-Darling tests. For determining significant firing rate modulation during hunting epochs (across epoch types and valence) across multiple sessions, a Friedman test with Tukey-Kramer post hoc (e.g. Figure 2G) or Cochran’s Q (i.e. Figure S2) was used. Correlation strength and significance between firing rates and behavioral state was calculated using Pearson’s *r*. To compare multiple groups to a fixed mean (usually 0), we used one-sample t-tests with Bonferroni correction. Statistical significance was considered to be p<0.05 and is indicated as * for p<0.05, ** for p<0.01, *** for p<0.001, and **** for p<0.0001.

### RESOURCE AVAILABILITY

#### Lead Contact

Further information and requests for reagents and resources should be directed to and will be fulfilled by the Lead Contact, Gina Turrigiano.

#### Materials Availability

This study did not generate new reagents.

#### Data and Code Availability

Processed data that went into generating figures will be deposited in a publicly available database such as Figshare upon acceptance. Large unprocessed datasets will be made available upon request. All code will be deposited at Github upon acceptance.

## ACKNOWLEDGEMENTS

We thank members of the Turrigiano lab for many helpful discussions, Lirong Wang and Zhe Meng for help with animal colony maintenance, the Brandeis Light Microscopy Core Facility, and the Brandeis Machine Shop for their assistance. XPRO1595 was a gift from INmuneBio (Boca Raton, FL, USA). Funding provided by T32NS007292 (BJL, NFW, MRS), R01 EY025613 (GGT), R35NS111562 (GGT), and Simons Foundation grant 901199 (GGT).

## AUTHOR CONTRIBUTIONS

Conceptualization: D.P.L, D.B., B.J.L., G.G.T.; Methodology: D.P.L., B.J.L., B.A.C., D.B., G.G.T.; Software: D.P.L., B.A.C., W.W., B.J.L., M.R.S., N.F.W.; Formal Analysis: D.P.L., B.A.C., D.B., W.W., M.R.S., N.F.W., K.B.B.; Investigation: D.P.L, D.B., W.W., B.J.L., M.R.S., N.F.W., K.B.B.; Writing: D.P.L., B.A.C., G.G.T.; Visualization: D.P.L., B.A.C., D.B., W.W., M.R.S., N.F.W.; Data Curation: D.P.L., D.B.; Supervision: G.G.T.; Funding Acquisition: G.G.T.

