## Supplemental Data for "Rapid prey capture learning slowly re-sets activity set points in rodent binocular visual cortex"

**Figure S1. Extended prey capture learning analyses of critical period rats.**

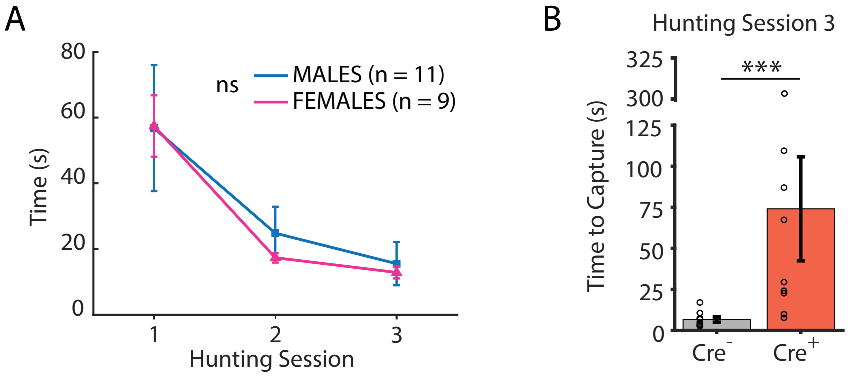


**Figure S1, related to Figure 1. Extended prey capture learning analyses of critical period rats. A.** Average time to capture, same population as in Fig. 1E, separated by sex. Repeated Measures Two-way ANOVA with Tukey correction, p=0.7. **B.** Average time to capture for PV-Cre- vs. PV-Cre+ rats during hunting session 3. Individual circles represent individual rats (Cre-: n=9, Cre+: n=9). Wilcoxon Rank Sum test: p=0.0005.

**Figure S2. Movement speed across hunting sessions, and modulation of firing rates in V1b by speed**

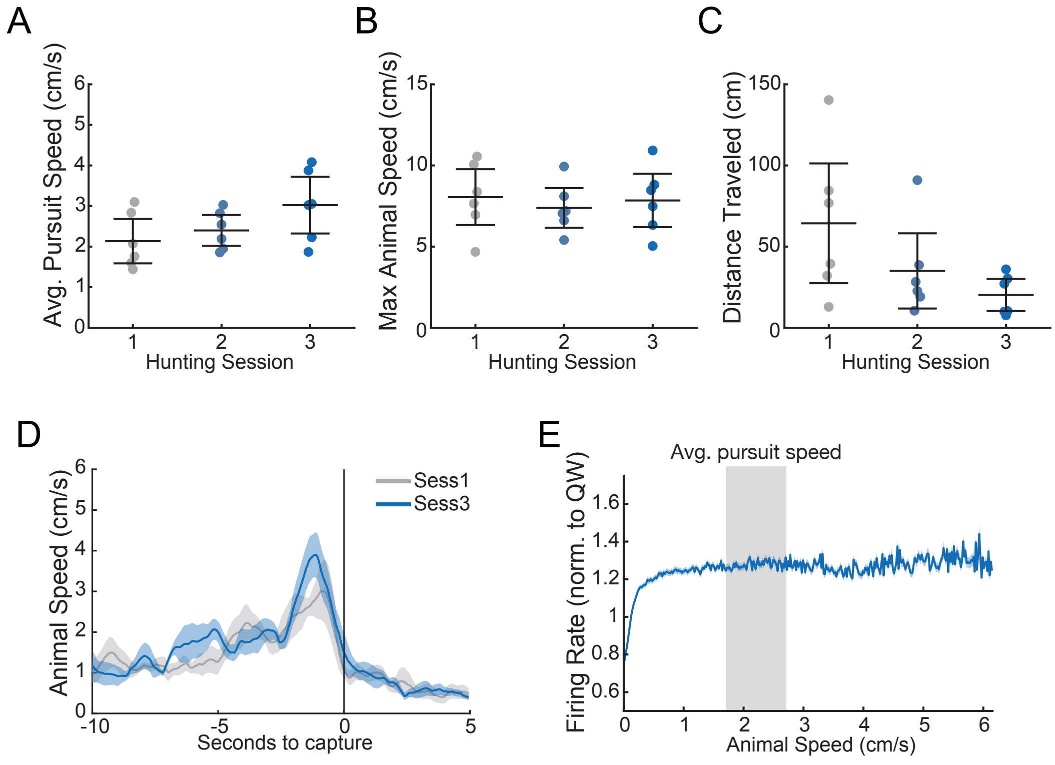


**Figure S2, related to Figure 2. Movement speed across hunting sessions, and modulation of firing rates in V1b by speed.** Average behavioral metrics across hunting sessions for chronically recorded animals in Figures 3-5 (n = 6 animals, each dot is single animal average). **A.** Average speed during pursuit for each animal. Repeated measures ANOVA (rmANOVA): p=0.09. **B.** Average max speed achieved during pursuit, rmANOVA: p=0.4. **C.** Average distance traveled during pursuit, rmANOVA: p=0.4. **D.** Average animal speed aligned to capture, session 1 versus 3. **E.** Animal movement speed vs. firing rate (normalized to quiet wake). Data from light period day of hunting. Grey bar indicates range of speeds during pursuit.

**Figure S3. Firing rate changes in V1b and V1m for sham and hunt conditions**

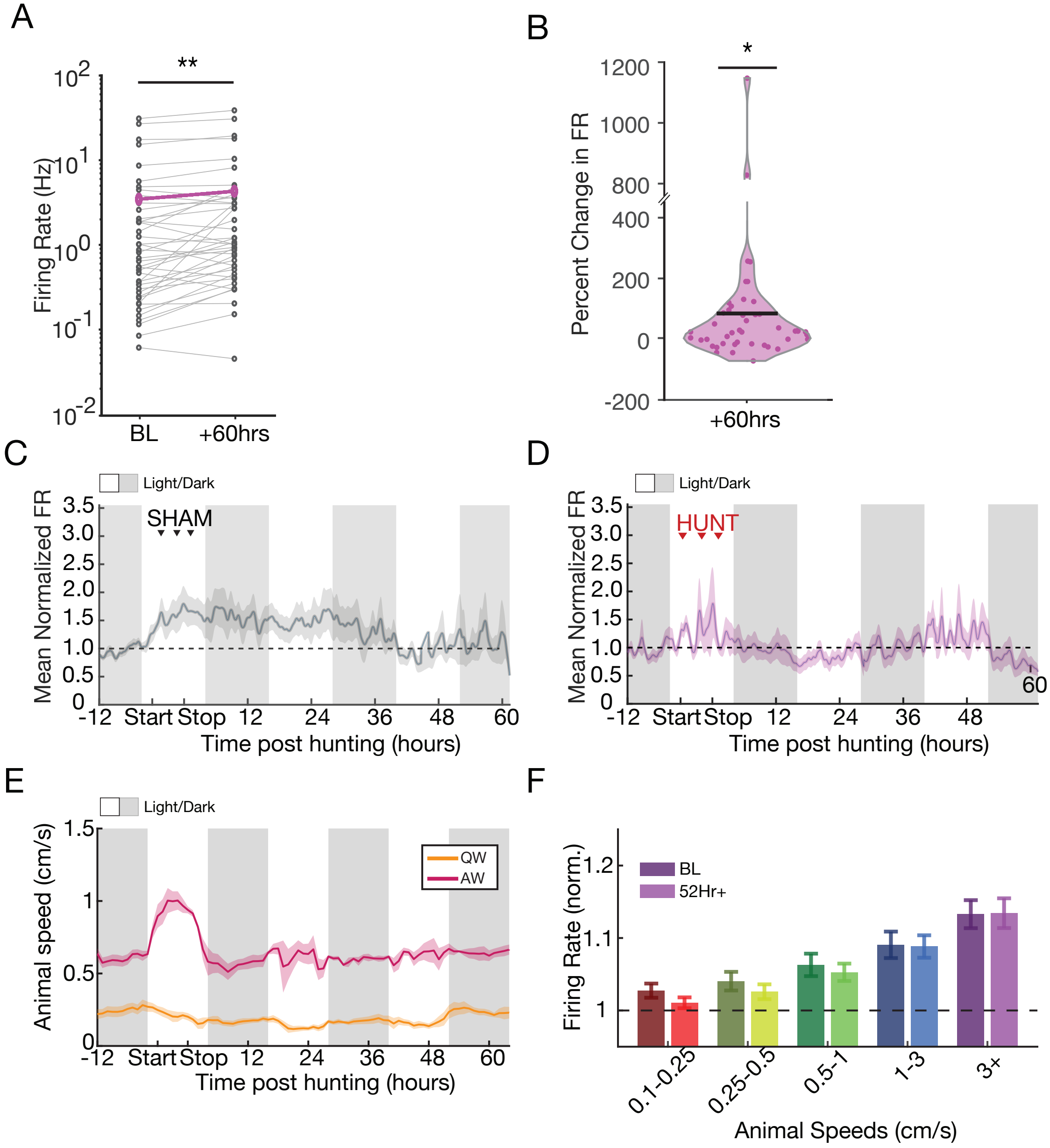


**Figure S3, related to Figure 3. Firing rate changes in V1b and V1m for sham and hunt conditions. A.** Ladder plot of RSU firing rates for all continuously recorded RSUs (n=45 neurons, individual dots; average indicated by magenta line) comparing baseline and 60 hours post-learning, Wilcoxon Sign Rank test, p=0.003. **B.** Distribution of percent change in RSU firing rates at 60 hours for same data in (A); black line denotes average.

One sample t-test, p=0.01. **C.** Mean normalized FRs of continuously recorded RSUs in the Sham condition (n=19 from 6 animals). **D.** Mean normalized FRs of continuously recorded RSUs from V1m (n=12 from 4 animals). **E.** Average animal speed for chronically-recorded hunt animals included in analysis for Fig 3 (n = 6), separated into active and quiet wake. **F.** Modulation of firing rate by speed for same population as in (E), comparing baseline dark period to dark period at 52 hours post hunt. For each bin and timepoint, firing rates for each neuron were normalized to those at near zero speeds and averaged. Two-way repeated measures ANOVA: animal speeds: p<0.0001, BL-52+ comparison: p=0.4, interaction: p<0.0001.

**Figure S4. Extended analyses of sleep/wake data**

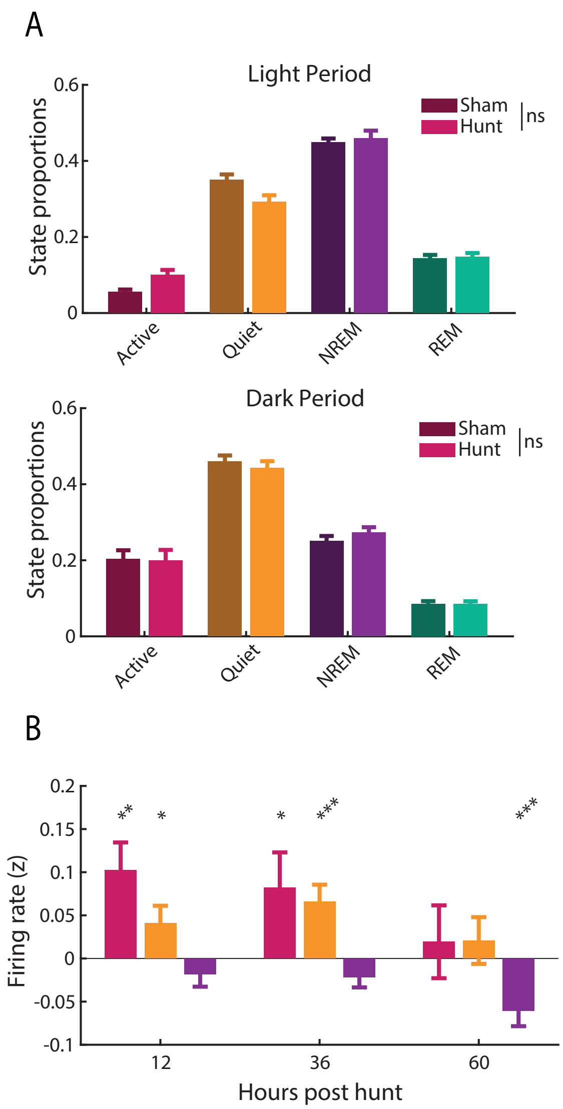


**Figure S4, related to Figure 5. Extended analysis of sleep/wake data. A.** Proportion of time spent in each state for Sham and Hunt animals during the light period **(top)** and dark period **(bottom)**. One-way MANOVA testing for effect of Sham vs. Hunt: light period, p=0.14, dark period, p=0.50. **B.** Firing rate change across extended wake/sleep episodes measured within Active/Quiet wake and NREM states in hunt animals. Calculated as z-scored change from within-state mean firing rates. Wilcoxon Sign Rank test with

Bonferroni corrected p-values: 12hr AW $p=0.004$, QW $p=0.03$, NREM $p=0.7$; 36hr AW $p=0.1$, QW $p=0.001$, NREM $p=0.08$; 60hr AW $p>0.9$, QW $p=0.4$, NREM $p=0.0002$.

**Figure S5. Extended mouse prey capture learning spine analysis**

A

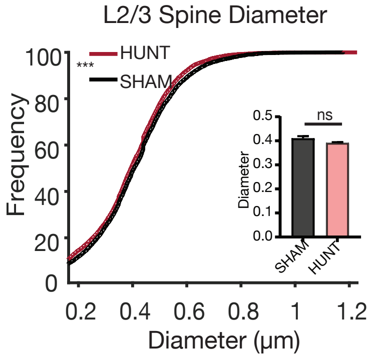


**Figure S5, related to Figure 6. Extended mouse prey capture learning spine analysis. A.** Cumulative distribution of L2/3 spine head diameter on day 6. Sham: n=5730 spines, Hunt: n=6831 spines, from 6 neurons and 3 animals per condition. Kolmogorov-Smirnov test, p <0.0001. Inset shows mean spine head diameter by neuron; Unpaired t-test, p=0.06.

**Figure S6. Active and passive properties of V1b pyramidal neurons measured ex vivo**

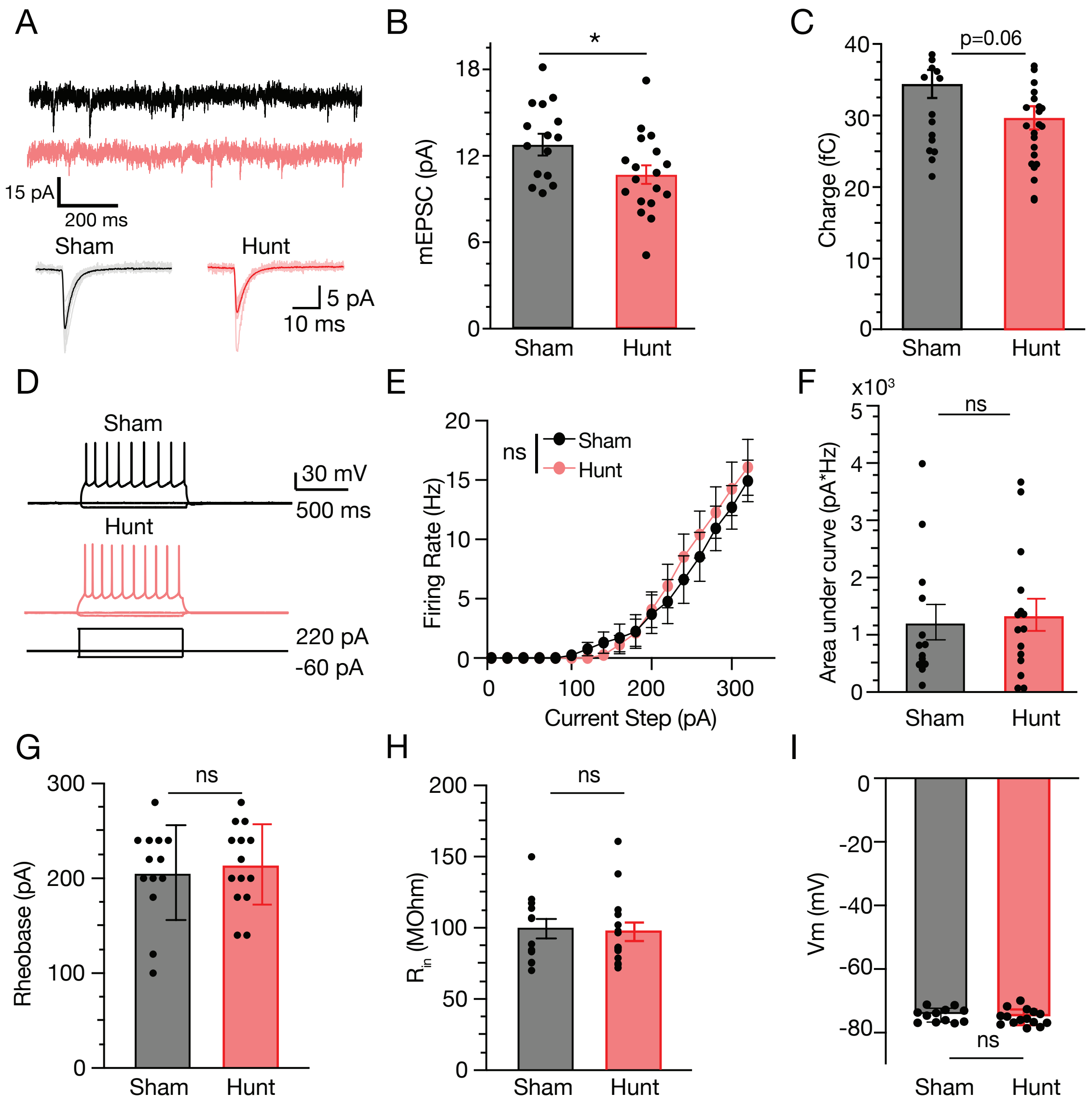


**Figure S6, related to Figures 6 and 7. Active and passive properties of V1b pyramidal neurons measured ex vivo. A.** Example mEPSC recordings and average waveforms (bottom) from example Sham (black) and Hunt (red) L2/3 pyramidal neuron. **B, C.** Average mEPSC amplitude (**B**) and charge (**C**) for Sham (grey) and Hunt (red) condition. Sham: 15 neurons from 5 animals. Hunting: 18 neurons from 5 animals. Wilcoxon Rank Sum test, p=0.01 for amplitude, p=0.07 for charge. **D.** Example whole cell current clamp recordings from Sham (grey) and Hunt (red) neurons in response to hyper- or depolarizing current injections. **E.** Frequency vs. Current (F-I) curve for Sham (black) and Hunt (red) conditions. Here and below: (**F-I**), Sham (grey): 13 neurons from 5 animals; Hunting (red): 15 neurons from 5 animals. Repeated Measures two-way ANOVA, p=0.8. **F.** Area under F-I curve, **G.** Rheobase, **H.** Input resistance, **I.** Resting potential; Wilcoxon Rank Sum tests (**F-I**), p=0.7, p=0.7, p=0.7, p=0.4.

**Movie S1.** Rat prey capture learning example from Hunting Session 1, related to Figure 1.

**Movie S2.** Rat prey capture learning example from Hunting Session 3, related to Figure 1.

**Movie S3.** PV-Cre negative rat (control) prey capture learning example from Hunting Session 3, related to Figure 1.

**Movie S4.** PV-Cre positive rat prey capture learning example from Hunting Session 3, related to Figure 1.
